# An extracellular vesicle delivery platform based on the PTTG1IP protein

**DOI:** 10.1101/2023.08.18.553853

**Authors:** Carla Martin Perez, Xiuming Liang, Dhanu Gupta, Mariana Conceição, Imre Mäger, Samir EL Andaloussi, Matthew J.A. Wood, Thomas C. Roberts

**Affiliations:** Department of Paediatrics, University of Oxford, John Radcliffe, Headington, Oxford, OX3 9DU, UK; Department of Physiology, Anatomy and Genetics, University of Oxford, South Parks Road, Oxford, OX1 3QX, UK; Biomolecular Medicine, Clinical Research Center, Department of Laboratory Medicine Karolinska Institutet, Stockholm, Sweden; Institute of Developmental and Regenerative Medicine, University of Oxford, IMS-Tetsuya Nakamura Building, Old Road Campus, Roosevelt Dr, Headington, Oxford OX3 7TY; MDUK Oxford Neuromuscular Centre, South Parks Road, Oxford, OX1 3QX, UK

## Abstract

Extracellular vesicles (EVs) hold great promise as therapeutic delivery vehicles, although their potential is limited by a lack of efficient EV engineering strategies to enhance loading and functional delivery of therapeutic cargo. Here, we identified N-glycosylation as a putative EV-sorting feature using a novel bioinformatics analysis strategy. We exploited this finding to develop a platform for EV-mediated delivery of macromolecular cargoes based on PTTG1IP, a small N-glycosylated, single spanning transmembrane protein. We demonstrate that PTTG1IP EV enrichment is dependent on its N-glycosylation at two sites, and that PTTG1IP is a suitable scaffold for EV loading of therapeutic cargoes. To achieve functional delivery, cargoes were fused to PTTG1IP, separated by various self-cleaving sequences intended to promote cargo release from the scaffold after EV loading. In addition, the fusogenic VSVG protein was co-expressed to enhance endosomal escape. This PTTG1IP-based strategy enabled highly efficient functional delivery of Cre protein to recipient cells and mouse xenograft tumors, as well as Cas9 and Cas9/sgRNA complex delivery to reporter cells. Importantly, PTTG1IP exhibited improved protein delivery potential relative to a scaffold based on CD63 (a common EV marker). Moreover, we developed various PTTG1IP variants with improved properties, demonstrating the versatility of PTTG1IP as an EV scaffold. The EV-loading platform described in this study offers significant advantages over other strategies, such as favourable membrane topology, the potential for further engineering, and functional delivery capability, which will enable the development of improved EV-based therapeutics.

## Introduction

Extracellular vesicles (EVs) are cell-derived, membranous nanoparticles released by cells into the extracellular environment that mediate cell-to-cell communication in both physiological and pathological conditions. EVs facilitate the intracellular transfer of proteins, lipids and nucleic acids^1^ to recipient cells and are able to regulate cellular signalling pathways by displaying specific molecules on their surface (e.g. membrane-bound ligands or receptors).

The unique properties of EVs as delivery vehicles (i.e. their biocompatibility, cargo versatility, and capacity for engineering) make them attractive candidates for therapeutic applications. However, several challenges must be addressed before the potential of EVs as therapeutics can be fully realized. Loading of therapeutic cargo molecules constitutes one of the main challenges for EV-based therapy. Not only must this cargo be present in the EVs, but it also needs to be capable of reaching the desired intracellular location once taken up by the recipient cells. Current methods for endogenous EV-protein loading often rely on tetraspanins such as CD63,^2–5^ that despite showing a high loading capacity are not readily amenable to further engineering on their extracellular domains (e.g. addition of targeting peptides).^6^ This is because that their N- and C-termini are both situated on the luminal side, and successful modification of the extraluminal loops can be challenging. There is therefore a need for more suitable EV scaffold proteins that allow for both highly efficient intracellular EV-loading and which can be modified with targeting moieties at the EV surface. The enrichment of specific proteins in EVs relative to expression levels in their respective producer cell suggests that there are mechanisms that regulate the sorting of protein cargo.^7^ Although these mechanisms are beginning to be deciphered, they are still not fully understood. Interestingly, protein features including post translational modifications (PTMs), short protein motifs, and protein domains have been shown to play a major role in EV-protein sorting and EV biogenesis.4 For example, the late domain motifs in viral Gag proteins can be recognized by the EV biogenesis machinery (i.e. ALIX and ESCRT-I component TSG101), which mediate their sorting and release in EVs.^9,10^ The CAAX motif found in Rab GTPases promotes their geranylgeranylation, which enables Rab binding to cellular membranes.^11^ In addition, palmitoylation and myristylation motifs have also been used extensively to promote protein EV-loading.^12–14^

We were motivated to identify additional PTMs (and other sequence features) associated with EV loading that may be exploited for therapeutic purposes. To this end, we developed a novel bioinformatics analysis: (IPFA, integrated proteomics and feature annotation analysis) that enabled the identification of EV-enriched annotated PTMs, protein motifs, and domains by combining protein feature annotations from UniProt with experimentally-observed EV-enriched proteins in proteomics data. This analysis identified several promising features, of which N-glycosylation was selected as the most promising PTM associated with EV protein enrichment. PTTG1IP (Pituitary tumor-transforming gene 1 protein-interacting protein), a single-pass transmembrane protein with two annotated N-glycosylation sites, was further investigated as the leading candidate EV-loading scaffold protein. PTTG1IP was found to be highly enriched in EVs in an N-glycosylation dependent manner, and loading of PTTG1IP-cargo fusion proteins was found to be highly efficient. Functional delivery of Cre recombinase to reporter cells and mouse xenograft tumors was accomplished with high efficiencies using PTTG1IP as an EV scaffold. In addition, the PTTG1IP platform was also utilized for successful Cas9 and Cas9/sgRNA complex delivery to reporter cells. Several PTTG1IP variants with improved properties for therapeutic delivery were also developed, demonstrating the potential for further engineering PTTG1IP as an EV scaffold beyond its native human amino acid sequence.

## Materials and methods

### Plasmids

Plasmids pSpCas9(BB)-2A-Puro (PX459) and pCMV-VSVG plasmid (pMD2.G) were kind gifts from Feng Zhang^15^ (Addgene plasmid #62988; https://www.addgene.org/62988/; RRID: Addgene_62988) and Didier Trono (Addgene plasmid #12259; https://www.addgene.org/12259/; RRID: Addgene_12259), respectively. All other constructs were cloned in-house by standard recombinant DNA technology techniques.

### Cell culture

Human embryonic kidney (HEK) 293T, T47D, HeLa, B16, and SKBR3 cells were cultured at 37°C with 5% CO2 in Dulbecco’s Modified Eagle Medium GlutaMAX (DMEM) (Gibco, Waltham, MA, USA) supplemented with 10% foetal bovine serum (FBS) and 1% Antibiotic-Antimycotic (PSA: Penicillin, Streptomycin and Amphotericin B) (Thermo Fisher Scientific, Waltham MA, USA).

T47D, HeLa, and B16 traffic light Cre reporter cells^16^ (30,000 cells per well) were seeded in 96- well plates. Traffic light cells constitutively express RFP, and upon Cre recombination also express GFP (when a floxed stop cassette placed before RFP is excised by Cre.). After 24h, cells were treated with EVs in 65 µl of complete Opti-MEM (Thermo Fisher Scientific). The media was changed to complete media and analyzed by flow cytometry 24h later.

HEK293T stoplight light Cas9 reporter cells^17^ (30,000 cells per well) were seeded in 96-well plates. For experiments using sgRNA expressing cells, sgRNA was transfected 24h before EV treatment. Cells were transfected with 45 ng of plasmid DNA/well using polyethylenimine (PEI) (Sigma-Aldrich, St. Louis, MO, USA) at a 4:1 ratio PEI:DNA in complete media. 24h later media was removed, and cells were treated with EVs in 65 µl of complete Opti-MEM. For non-sgRNA expressing experiments, cells were treated with EVs in 65 µl of complete Opti-MEM (Thermo Fisher Scientific). In both cases, media was changed to complete media after 24h and reporter cell activation was analysed by flow cytometry.

### EV production

For cell culture experiments, at least four 150 mm plates of HEK293T cells per condition were transfected with 35 µg DNA/plate. For *in vivo* experiments, sixty 150 mm plates of HEK293T cells per condition were transfected with 35 µg DNA/plate. 24h later, the media was replaced with Opti-MEM and after a further 48h, cell culture supernatant was collected for subsequent EV isolation.

### EV isolation

Cell culture supernatant was collected and centrifuged consecutively at 500 *g* and 2,000 *g* to remove cell debris and large particles, respectively. The media was then filtered through a 0.45 µm filter (STARLAB, Hamburg, Germany) and concentrated using a 10 KDa Vivaflow tangential flow filtration (TFF) membrane (Sartorius, Göttingen, Germany) to a volume of 10-15 ml. Media was concentrated further to a final volume of 1-2 ml using 10 KDa Amicon spin filters (Merck Millipore, MA, USA) by centrifugation at 3,500 *g*. Concentrated media was then subjected to size exclusion chromatography using Sepharose 4 fast flow column connected to the ÄKTA prime system (both GE Healthcare, Chicago, IL, USA) and eluted in PBS with a flow rate of 0.5 ml/min. EV and protein fractions were selected based on 280 nm absorbance. The yield of vesicles was determined using single-vesicle analysis.

### Single vesicle analysis

Single vesicle analysis was performed to characterize isolated EVs and conditioned media. For isolated EVs, samples were analysed directly. For conditioned media analysis, HEK293T cells were seeded on 12-well plates (100,000 cells per well) 24h before transfection. The following day, cells were transfected with 700 ng of plasmid DNA/well using polyethylenimine (PEI) (Sigma-Aldrich, St. Louis, MO, US) at a 4:1 ratio PEI:DNA in Opti-MEM (Thermo Fisher Scientific). 24h later, media was replaced with Opti-MEM and 48h after media replacement, conditioned media was collected. Before single-vesicle analysis, condition media was centrifuged first at 500 *g* to remove cells and subsequently at 2,000 *g* to remove large particles. After 0.22 µM filtering (STARLAB), EVs or conditioned media were analysed on a Flow NanoAnalyzer U30 instrument (NanoFCM, Nottingham, UK). 2,000-12,000 single particle events were counted for 1 min using light scattering and 488/24 nm blue laser set to 10 mW and 10% SS decay, at a sampling pressure of 1.0 kPa. Data were analysed using NanoFCM Professional Suite v1.8 software (NanoFCM). EV size and concentration were determined by interpolation from a standard curve of silica nanoparticles with diameters ranging from 68 nm to 155 nm. Non-transfected HEK293T cell conditioned media was used as a fluorescence negative control to set the threshold for GFP positivity.

### Flow cytometry

Cells were washed with PBS followed by 15 min incubation with 50 µl TrypLE Express (Thermo Fisher Scientific). Trypsin was inactivated using 100 µl FACS buffer (PBS containing 10% FBS) and cells were kept on ice until ready for analysis. Cells were analysed on a CytoFLEX LX flow cytometer (Beckman Coulter, Brea, CA, USA) and data analysis was performed using FlowJo software (BD Biosciences, Franklin Lakes, NJ, US). Activated reporter cells were identified according to their GFP expression (measured at 488-525 nm).

### N-glycosylation assays

PNGase F and Endonuclease H treatments were performed to assess the N-glycosylation status of PTTG1IP. For both treatments, 20 µg protein lysate from HEK293T cells transfected with the experimental plasmid constructs was denatured for 10 min at 100°C with 10× Glycoprotein Denaturing Buffer. For PNGase F, this was followed by incubation with 1 µl PNGase F (NEB, Ipswich, MA, USA), 2 µl 10% NP-40 (NEB) and 2µl 10× GlycoBuffer 2 (NEB) buffer at 37°C 1h. For Endonuclease H treatment, after denaturation, samples were incubated with 1.5 µl EndoH and 2 µl 10× GlycoBuffer 3 (all NEB) buffer at 37°C 1h. Glycosylation status was subsequently inferred by assessing shifts in protein size by Western blot.

### Western blot

Cells were lysed in 1× RIPA buffer (Thermo Fisher Scientific) containing 1× cOmplete protease inhibitor cocktail (Merck). Protein from EV samples was concentrated 10× by acetone precipitation whereby four volumes of cold (-20°C) acetone was added and samples were incubated for 24h at -20°C. After precipitation, samples were centrifuged at 5,000 *g* for 20 min. The resulting pellet was resuspended in 1× RIPA buffer containing 1× cOmplete protease inhibitor cocktail and 1x NuPage sample reducing agent (Thermo Fisher Scientific). Protein concentration was determined by Bradford assay (Sigma-Aldrich) following manufacturer’s instructions. 5-10 µg of total protein was heated at 70°C for 10 min and then resolved on a 4–12% NuPAGE gel (Thermo Fisher Scientific) by SDS-PAGE. Gels were run at 150 V for 2h in 1× NuPAGE Tris-Acetate SDS Running Buffer (Thermo Fisher Scientific). Protein was transferred onto a PVDF membrane (Merck) at 60 V for 90 min in 1× NuPAGE Transfer Buffer (Thermo Fisher Scientific). Membranes were blocked in Intercept (PBS) Blocking Buffer (Sigma-Aldrich) for 30 mins at room temperature and incubated in Blocking Buffer with primary antibodies at 4°C overnight. Blots were washed with PBS supplemented with 0.1% Tween-20 (PBS-T) three times and incubated with secondary antibody for 1h at room temperature. Antibody information is described in **Supplementary Table 1**. Blots were imaged using a LI-COR Biosciences Odyssey Fc instrument (LI-COR, Lincoln, NE, USA).

### *In vivo* EV-transfer

Animal experimentation was conducted at Karolinska Institutet with institutional ethical approval (permit 2173-2021). Mice (C57BL/6, female, 9 weeks old) were injected with 5×10^5^ B16F10 Cre reporter cells/mouse 10 days before EV treatment in order to generate xenograft tumors. On day 10, isolated EVs were administered to recipient mice by intratumoral injection (1×10^10^ EVs/mouse). 4 days later, tumors were harvested and Cre activity was analyzed by immunohistochemistry (IHC) and PCR. For IHC, half of the tumor tissues were fixed with paraformaldehyde (PFA) for generating slides. For PCR, the other half of tumor tissues were lysed using lysis buffer (0.1% Triton X-100) and a TissueLyser homogenizer (Qiagen, Hilden, Germany). DNA for PCR was isolated from 50 µl of tissue lysate using the Maxwell RSC tissue DNA Kit (Promega, Madison, WI, USA).

### Cre-mediated LoxP site recombination analysis

To PCR amplify the LoxP-Cre recombination site in B16 Cre reporter cells, forward primer 5ʹ- CGCAAATGGGCGGTAGGCGTG-3ʹ and reverse primer 5ʹ-GTGGTGCAGATGAACTTCAGGGT-3ʹ were used. The forward primer was located in the CMV promoter, before the first LoxP site, while the reverse primer located after the second LoxP site at the 5ʹ end of the GFP reporter. PCR was performed using genomic DNA isolated from tumors as the template using HotStarTaq Plus Master Mix Kit (Qiagen) according to the manufacturer’s protocol. PCR products were analyzed on a 1% agarose gel using VersaDoc 3000 Gel Imaging System (Bio-Rad, Hercules, CA, USA). The intensity of the bands was quantified by densitometric analysis using ImageJ 1.53k software (NIH, Bethesda, MD, USA).

### Immunochemistry of tumor xenografts

The tumor xenograft tissue slides were fixed at 65°C for 1 hour. Deparaffinization and rehydration were performed by incubating the slides in several solvents as follows: xylene 20 min, 100% ethanol 3 min twice, 95% ethanol 3 min, 70% ethanol 3 min, and 50% ethanol 3 min. Slides were subsequently rinsed with cold running tap water for 5 min followed by antigen retrieval using citrate buffer at pH 6.0 (Sigma Aldrich). Afterwards, slides were washed 3 times with PBS (5 min each), and then slides were blocked using blocking buffer for 30 min at 37°C. Slides were then incubated overnight with anti-GFP antibody (1:200 dilution, ab290, Abcam, Cambridge, UK). The next day, after 3 washes in PBS, slides were incubated with secondary antibody Goat Anti Rabbit IgG H&L (Alexa Fluor 488) (1:500 dilution, Abcam, ab150077) for 30 min at 37°C. After 3 washes with PBS, slides were mounted using ProLong Diamond Antifade Mountant with DAPI (Thermo Scientific) and images were acquired using a confocal microscope (Nikon, Tokyo, Japan).

### Nanoluciferase assay

EVs were harvested and recipient cells treated as described above. Doses of EVs were determined according to the input luminescence of the preparations. After incubation, the cells were washed twice with PBS and lysed in 100 µl of PBS with 0.1% Triton X-100. The plate was then incubated on an orbital shaker at room temperature for 10 min for complete lysis of the cells. The cell lysate was then analyzed for nanoluciferase activity using the appropriate substrates as described previously.^18^

### Statistics and bioinformatics

GraphPad Prism software version 9 (GraphPad, San Diego, CA, USA) was used to perform statistical analysis and generate graphs. Differences between two groups were assessed by Student’s *t*-test. Differences between more than two groups were tested by one-way ANOVA and Bonferroni *post hoc* test.

The integrated proteomics and feature annotation analysis (IPFA) was performed using custom Python (version 3.8) scripts with the following dependencies: *numpy*, *seaborn*, *functools*, *math*, *matplotlib*, *scipy*, *json*, and *request*. Differences between the distributions of EV:cell abundance ratios for each feature annotation category relative to the distribution for all proteins were tested by Kolmogorov-Smirnov test followed by Benjamini-Yekutieli *post hoc* test. In order to deal with the missing data problem, a value of ‘one’ was added to every mass spectrometry area measurement (abundance). As such, the mathematical distances between samples were maintained and any ‘zeros’ removed.

## Results

### *In silico* identification of EV-loading features

Protein features including post-translational modifications (PTMs), protein motifs, and protein domains (such as ubiquitin-like modifications)^19^ have been shown to play a role in EV-protein sorting.^8^ Therefore, to identify novel features that could lead to protein sorting to EVs we developed a bioinformatics approach; IPFA (integrated proteomics and feature annotation analysis). IPFA uses custom Python scripts to combine mass spectrometry proteomics data (from matched EVs and producer cells) with PTM, motif, and domain feature annotation data derived from UniProt (**Supplementary Fig 1**). The first step of the analysis was the extraction of feature annotations from the UniProt database.^20^ For PTMs and motifs, data were manually curated to remove duplicate features. Moreover, groups of related PTMs (e.g. glycosylation) were pooled and included as additional features. Data from protein features were then integrated with publicly available mass spectrometry proteomics data^21^ and features that were annotated on fewer than 5 proteins filtered out for reasons of statistical power. EV-enriched features were identified as those that were associated with significantly enriched proteins (relative to all proteins) by Kolmogorov-Smirnov test, and which exhibited an increase in the EV vs cell abundance ratio relative to the median ratio for all proteins (**Fig 1**).

**Figure 1.**
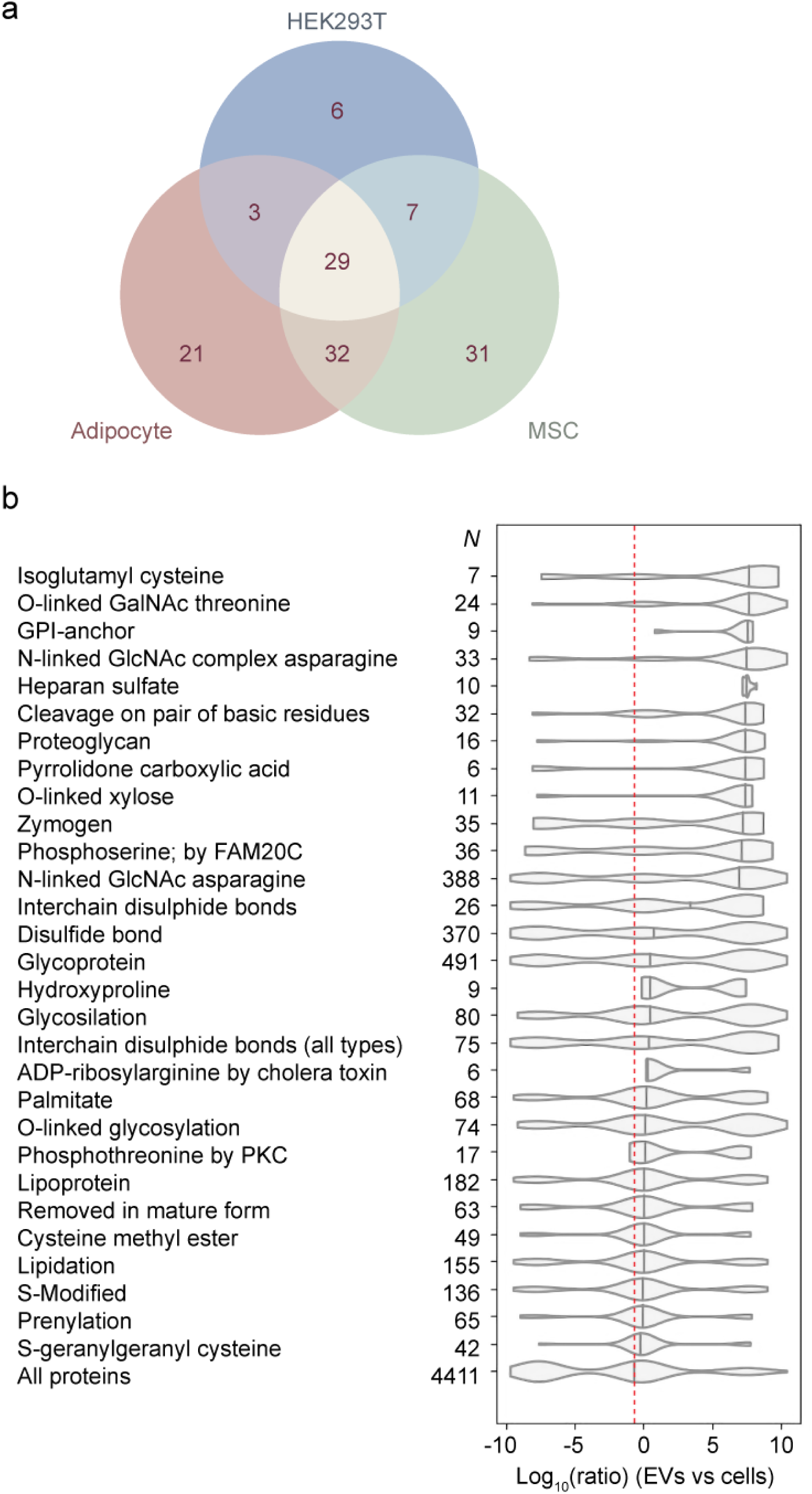
EV-enriched PTM annotations identified with IPFA analysis. To identify EV-enriched PTM annotations, IPFA analysis was performed for HEK293T, MSC and adipocyte datasets. (**a**) Venn diagram of statistically significant EV-enriched PTMs in HEK293T cells, MSCs, and adipocytes. Numbers in the diagram refer to the number of PTMs identified for each group. Violin plots representing the EV to cell abundance ratios (log10) of proteins with statistically significant EV-enriched PTM annotations identified in HEK293T cells (**b**). Only EV-enriched PTM annotations common across all three cell lines are shown. Numbers on the y-axis depict the number of proteins with the given PTM in the dataset. Violin plot of the EV to cell abundance ratios (log10) of all proteins in the dataset is also shown. Median value of all proteins in the dataset is represented as a dashed line.

To identify protein EV-enriched PTMs, we performed IPFA analysis in three different cell lines: HEK293T cells, mesenchymal stem cells (MSCs), and adipocytes. 53, 39, and 75 EV-enriched PTMs were identified in HEK293T cells, MSCs, and adipocytes, respectively (**Fig 1a**). Among the identified PTMs, 29 were common across all 3 cell lines (**Fig 1a**). Notably, N-linked GlcNAc asparagine (N-GlcNAc) was highly enriched and present in ∼10% of all proteins in the three datasets (**Fig 1b**, **Supplementary Fig 2a**,b). Lipidation was also present on numerous proteins (∼5%). Hydroxyproline, pyrrolidone carboxylic acid, and kinase specific phosphorylation were also identified as EV-enriched in all cell lines analysed.

EV-enriched protein motifs (**Supplementary Fig 3**) and domains (**Supplementary Fig 4**) were analysed in parallel. Among the identified EV-enriched motifs and domains the PTAP motif, SPRY domain, and MIT domain were selected for further analysis due to their potential applicability as EV-loading scaffolds.

### PTTG1IP enables highly efficient EV-loading in a N-glycosylation dependent manner

Selected candidate features (N-linked glycosylation, PTAP motif, SPYR domain, and MIT domain) were assessed for EV-loading by fusing these features to GFP and analysing their sorting to EVs by Western blot and single vesicle analysis (**Supplementary Fig 5a**). In the case of N-linked glycosylation and the MIT domain, we decided to test the smallest proteins containing these feature that could be found in our dataset as we reasoned that this would simplify the required protein engineering. PTTG1 Interacting Protein (*PTTG1IP*) and MIT domain-containing protein 1 (*MITD1*) were selected for N-glycosylation features and MIT domain, respectively (GFP was fused to the C-terminus of these proteins in both cases). Importantly, the PTTG1IP sequence also comprises a PSI domain (domain found in Plexins, Semaphorins, and Integrins) which was similarly found to be EV-enriched by IPFA analysis (**Supplementary Fig 4**). For the PTAP motif and SPRY domain, these protein features were fused directly to GFP (PTAP was fused to the C-terminus of GFP while SPRY domain was fused on either the C- or the N-terminus).

GFP was highly enriched in the EVs when fused to PTTG1IP (∼75% of GFP^+^ vesicles) as compared to GFP alone (i.e. no scaffold) (∼3% of GFP^+^ vesicles) (**Supplementary Fig 5** **b,d**). In addition, the mean fluorescence intensity of PTTG1IP-GFP vesicles was significantly higher (∼58%) than that of GFP alone, indicating a greater degree of GFP loading per vesicle (**Supplementary Fig 5c**). Fusion of the SPRY domain to GFP also increased GFP EV-loading, although to a much lesser extent than that observed for PTTG1IP (∼23% of GFP^+^ vesicles).

N-glycosylation sites annotated in UniProt include both experimentally validated and inferred sites according to sequence analysis. PTTG1IP contains two putative N-glycosylation sites on asparagines 45 and 54. Thus, to confirm the N-glycosylation status of PTTG1IP, we mutated residues N45 (N45Q), N54 (N54Q) or both N45 and N54 (2NQ) to glutamine in order to remove these potential N-glycosylation sites and subjected HEK293T lysates from cells transfected with PTTG1IP mutants to PNGase F (Peptide N-Glycosidase F) and Endonuclease H (EndoH) treatment (**Fig 2a**). PNGase F cleaves all N-glycosylation sites and EndoH removes only those glycosylation modifications of the high mannose or hybrid types.^22^ Treatment with PNGase F resulted in a shift in size for the wild-type (WT) protein from ∼55 kDa to ∼48 kDa, indicating that PTTG1IP is indeed glycosylated (**Fig 2b**). Single N-glycosylation mutants also exhibited higher molecular weights than for the 2NQ double mutant (**Fig 2b**). Taken together, the shifts in molecular weight of the three mutants as compared to the WT suggest that both sites are N-glycosylated. In contrast, after treatment with EndoH, a band shift was only observed for PTTG1IP N45Q, suggesting that N54 N-glycosylation is of high mannose or hybrid type while N45 is of the complex type (**Fig 2b**).

**Figure 2.**
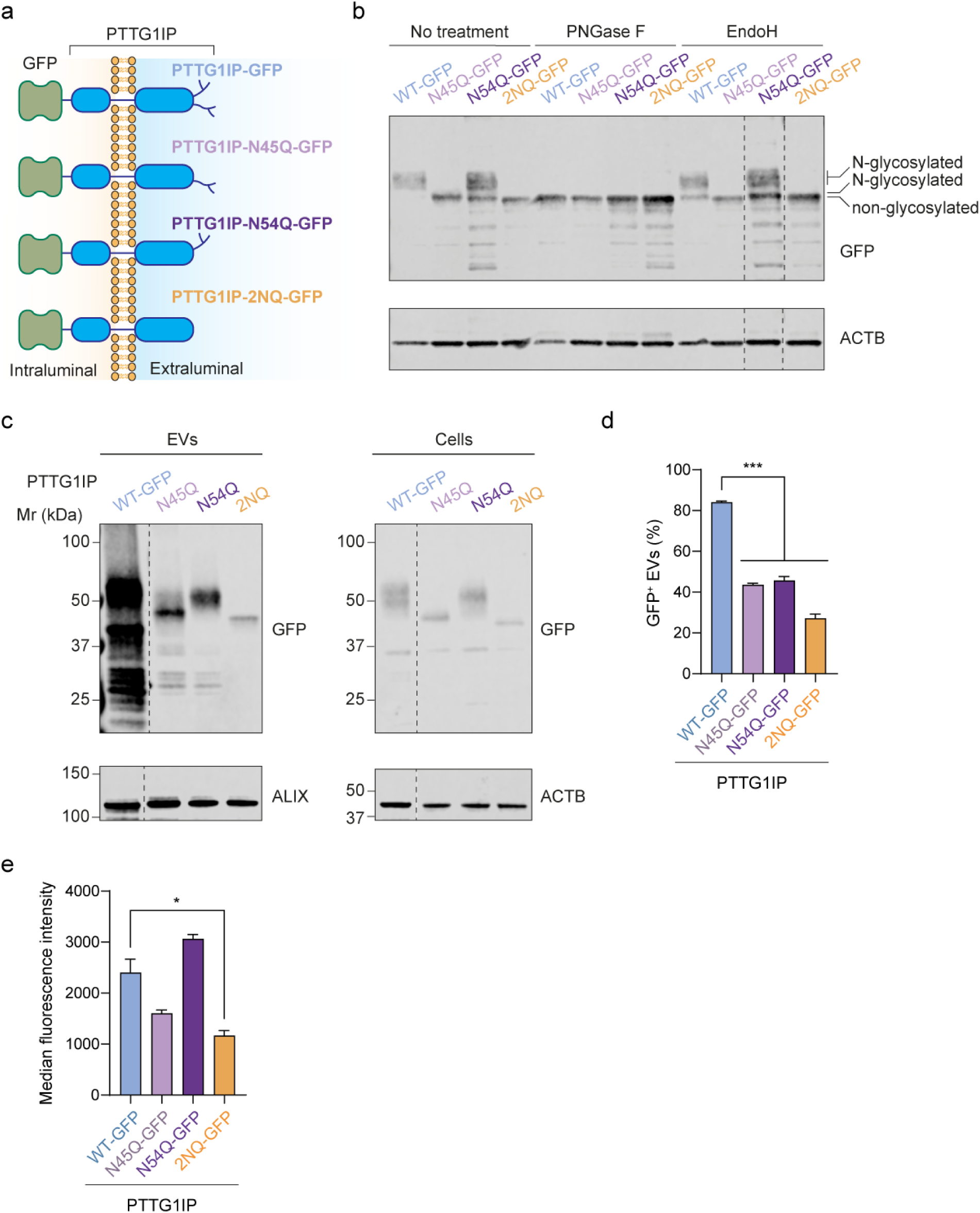
N-glycosylation of PTTG1IP promotes its sorting to EVs. (**a**) Scheme of PTTG1IP N-glycosylation mutants fused to GFP. (**b**) Western blot of WT PTTG1IP and NGlcNAc mutants fused to GFP in HEK293T cells. Cell lysates were treated with PNGase F to remove all N-glycosylation or with Endonuclease H (EndoH) to remove high-mannose and hybrid N-glycosylations. Lysates were blotted against ACTB and GFP. (**c**) Western blot of WT PTTG1IP and N-glycosylation mutants fused to GFP in HEK293T EVs and cells. Cell lysates and isolated EVs were blotted against ACTB and ALIX respectively. Both cell lysates and isolated EVs were blotted against GFP. (**d**) Percentage of GFP^+^ EVs quantified by single-vesicle analysis of conditioned media from HEK293T cells transfected with GFP-fusion constructs. (**e**) Median fluorescence intensity of GFP^+^ EVs. Statistical significance was tested by one-way ANOVA and Bonferroni *post hoc* test. Data are presented as mean + SEM, *n* = 3 replicates, \**P* < 0.05, \*\**P* < 0.01, \*\*\**P* < 0.001. Blots were modified as follows (indicated with dotted lines): In (**b**) the samples originate from the same blot and the order of the last two lanes was changed for consistency with other figures. In (**c**) the blot was modified to remove a lane that is irrelevant to the present manuscript. (Full uncropped blots available on request).

To determine if PTTG1IP EV-sorting was influenced by its N-glycosylation status, we assessed the EV-loading potential of PTTG1IP-GFP N-glycosylation mutants in HEK293T by Western blot and single vesicle analysis. PTTG1IP 2NQ-GFP EV-loading was markedly reduced when compared to WT (27.2% of GFP^+^ vesicles vs 84%) (**Fig 2c,d,e**). Mean fluorescence intensity of 2NQ mutant was also 48% that of WT (**Fig 2e**), which indicates that 2NQ GFP^+^ vesicles contain around half the amount of GFP per vesicle. The N45Q and N54Q single mutants also showed reduced loading (43.6% and 45.7% of GFP^+^ vesicles, respectively) as compared to WT PTTG1IP. Importantly, the expression levels of PTTG1IP mutant constructs in cells were similar to that of WT PTTG1P, indicating that the lower sorting to EVs is not a consequence of diminished expression of the loading scaffold (**Fig 2c**).

### PTTG1IP can be further engineered to improve its applicability as an EV-scaffold

PTTG1IP is a 180 amino acid single-pass type I transmembrane protein. It is composed of an N-terminal signal peptide and a PSI domain harbouring the two N-glycosylation sites in its extracellular domain, a transmembrane domain, and an intracellular C-terminus consisting of an unstructured domain followed by a coiled coil region and two endocytosis signals (**Fig 3a**). To improve the suitability of PTTG1IP as a scaffold for cargo loading to the EVs, we engineered two novel PTTG1IP variants. In the first variant, PTTG1IP-2YA, the two endocytosis signals of PTTG1IP (i.e. YXXL motifs)^23^ at the C-terminus were mutated to YXXA (**Fig 3b**). It has been previously shown that disrupting the endocytosis signal (YXXΦ motif, where Φ is a hydrophobic residue: L, I, M, F, V) of CD63 re-directs the protein towards the microvesicle pathway rather than to the MVB exosome pathway.^24^ In the second variant, PTTG1IP-Δ130-164, the coiled coil region of PTTG1IP (130 to 164 amino acids, in the intracellular region) was deleted (**Fig 3b**), as a sequence in this region has previously been proposed to interact with p53.^25^ To determine the EV-loading capacity of the variants, we transfected HEK293T with WT PTTG1IP-GFP, PTTG1IP- 2YA-GFP, or PTTG1IP-Δ130-164-GFP and assessed their loading into EVs by single vesicle analysis. Expression of both novel variants resulted in a similar percentage of GFP^+^ particles (∼80%) (**Fig 3c**). Nevertheless, the EV-mean fluorescence value for the PTTG1IP-2YA mutant was approximately twice the value of WT PTTG1IP, indicating that PTTG1IP-2YA was capable of loading double the amount of GFP per vesicle (**Fig 3d**). Taken together, these results show that the PTTG1IP sequence is amenable to further protein engineering.

**Figure 3.**
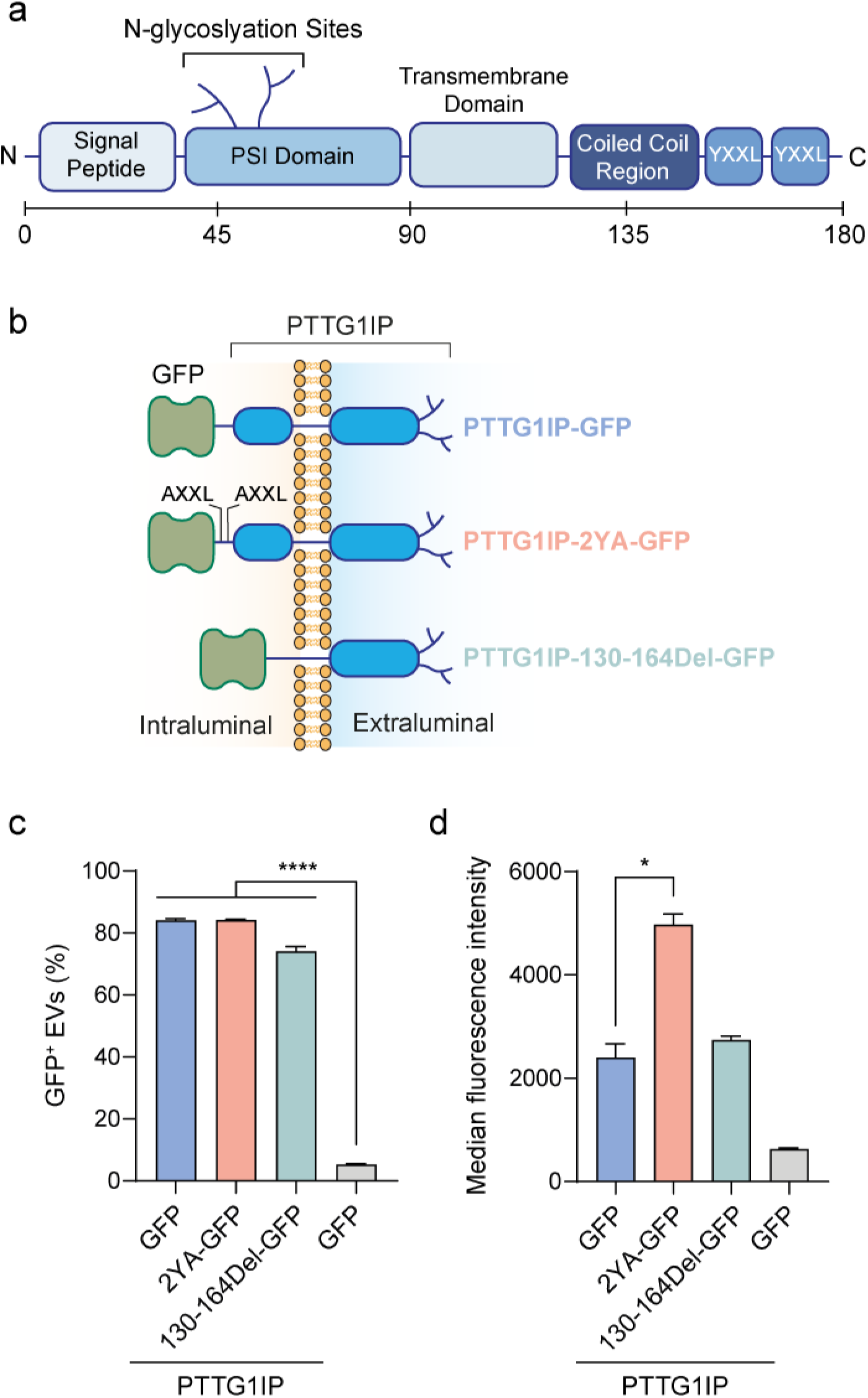
Loading of engineered PTTG1IP scaffolds into EVs is highly efficient. (**a**) PTTG1IP protein structure and length. (**b**) Scheme of PTTG1IP variants fused to GFP. PTTG1IP-2YA has its two YXXL endocytosis motifs mutated to AXXL and PTTG1IP-Δ130-164 has a deletion from amino acid 130 to164. (**c**) Percentage of GFP^+^ EVs quantified by single-vesicle analysis of conditioned media from HEK293T cells transfected with GFP-fusion constructs. (**d**) Median fluorescence intensity of GFP^+^ EVs. Statistical significance was tested by one-way ANOVA and Bonferroni *post hoc* test. Data are presented as mean + SEM, *n* = 3 replicates, \**P* < 0.05, \*\*\*\**P* < 0.0001.

### Assessment of Cre recombinase delivery to reporter cells

To investigate the ability of PTTG1IP to functionally deliver protein cargo to recipient cells, we assessed the transfer of Cre recombinase to reporter cells using a variety of fusion constructs with PTTG1IP as the loading scaffold (**Fig 4**). To achieve functional delivery in recipient cells, several obstacles need to be overcome. Firstly, the EVs need to be loaded with sufficient protein cargo in the producer cells. Secondly, the cargo needs to be free to reach its target intracellular location following uptake by the recipient cell. Lastly, once taken up by the recipient cells (in most cases through the endocytic pathway), the EVs must escape endosomes before they are either re-exported to the extracellular space or degraded by the lysosome. To address cargo-release from PTTG1IP, we utilized two different self-cleaving sequences (i) an intein derived from *Mycobacterium tuberculosis*^26,27^ (hereafter ‘intein’), and (ii) a sequence derived from *Shewanella oneidensis*^28^ (**Fig 4a**). Ideally, these sequences should self-cleave at lower pH so that cargo release only takes place in the intraluminal vesicles (in MVBs) of EV producer cells or in the late endosome of recipient cells. To promote endosomal escape, we co-expressed VSVG (vesicular stomatitis virus glycoprotein), a viral protein that mediates fusion of the EV and endosome membranes at lower pH therefore promoting endosomal escape.^29^ Importantly, VSVG and the cargo protein must be present on the same EVs to achieve functional delivery.

**Figure 4.**
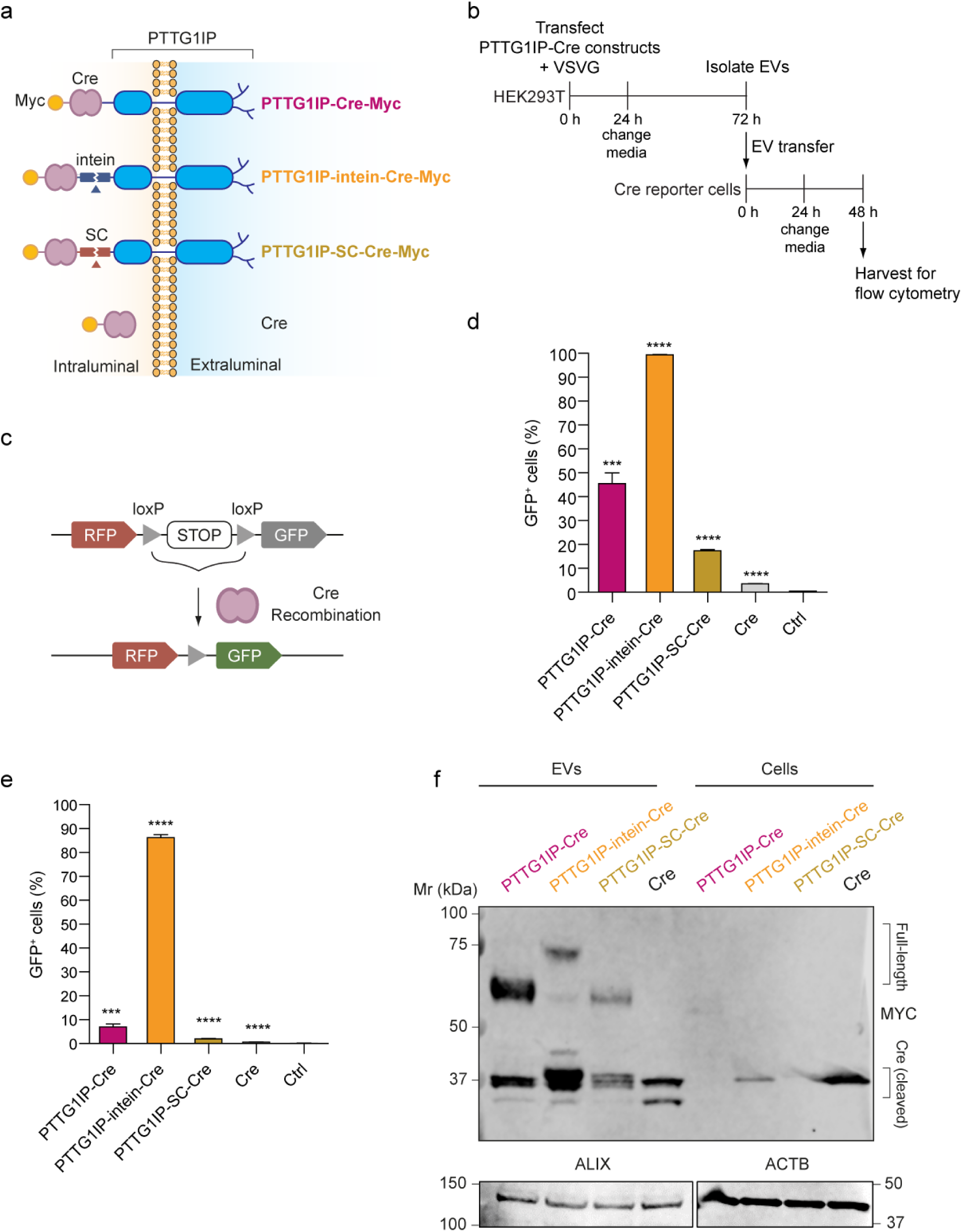
PTTG1IP-EV-mediated delivery of Cre recombinase to reporter cells. HEK293T cells were transfected with various PTTG1IP constructs and the capacity to transfer Cre protein was tested in reporter cells. (**a**) Scheme of tested Cre fusion constructs. SC is a self- cleaving endopeptidase from *Shewanella oneidensis*. The intein is a self-cleaving sequence from *Mycobacterium tuberculosis*. (**b**) Timeline of EV-transfer experiment to Cre reporter cells. 24h after donor cell transfection with Cre constructs and VSVG, media is replaced with fresh Opti- MEM and EVs are isolated 48h later by SEC. Isolated EVs are then transferred to recipient cells in 96-well plates (5×10^9^ EVs/ml). 24h later media is replaced with DMEM 10% FBS and after 24h cells are analysed for GFP expression by flow cytometry. (**c**) Scheme of the Cre reporter cells. RFP is constitutively expressed by reporter cells while GFP is only expressed when the floxed stop cassette is excised by Cre. (**d**) GFP positive cells quantified by flow cytometry 48h after EV- transfer to T47D cells. (**e**) GFP positive cells quantified by flow cytometry 48h after EV-transfer to HeLa cells. (**f**) Western blot of Cre-fusion constructs in HEK293T EVs and cells. Both cell lysates and isolated EVs were blotted against MYC. Cell lysates and isolated EVs were blotted against ACTB and ALIX, respectively. Statistical significance was tested by one-way ANOVA and Bonferroni *post hoc* test. Data are presented as mean + SEM, *n* = 3 replicates, \*\*\**P* < 0.001, \*\*\*\**P* < 0.0001.

To test delivery of Cre to recipient cells, we first transfected several Cre constructs (**Fig 4a**) together with VSVG into HEK293T producer cells and the resulting EVs were transferred to Cre reporter cells (**Fig 4b**). These reporter cells constitutively express RFP and contain a floxed stop signal followed by an in-frame GFP cassette. Upon Cre delivery, the LoxP sites recombine to excise the stop signal and GFP is expressed (**Fig 4c**). Constructs comprised of PTTG1IP fused to Cre (PTTG1IP-Cre), PTTG1IP fused to Cre separated by an intein (PTTG1IP-intein-Cre) or the self-cleaving sequence from *Shewanella oneidensis* (PTTG1IP-SC-Cre), and Cre alone. T47D or HeLa reporter cells were treated with 5×10^9^ EVs/ml and reporter activation was assessed 48h later by flow cytometry to determine the percentage of GFP positive cells. PTTG1IP-intein-Cre showed the highest activation (∼98% in T47D and ∼85% in HeLa cells), while PTTG1IP-Cre induced activation of ∼40% in T47D and ∼10% in HeLa cells (**Fig 4d,e**). Conversely, for Cre alone (no scaffold), activation was ∼3% in T47D and ∼1% in HeLa cells (**Fig 4d,e**). Notably, Cre was released in the EVs from PTTG1IP-Cre fusion protein without the need for a self-cleaving protein as observed from the ∼38 kDa band that corresponds to the expected size of Cre (**Fig 4f**), suggesting the presence of an intrinsic cleavage site contained within the PTTG1IP sequence itself. Nevertheless, when the intein is placed between PTTG1IP and Cre (PTTG1IP-intein-Cre) there was an increase of released Cre in the EVs (**Fig 4f**). By contrast, the other self-cleaving sequence derived from *Shewanella oneidensis* (PTTG1IP-SC-Cre) resulted in fainter Cre bands, indicating that SC was not as effective as the intein sequence for Cre release.

Inspection of the Cre Western Blot in **Fig 4f**, showed that free Cre and Cre released from the PTTG1IP fusion proteins were of similar sizes. We therefore reasoned that the site of intrinsic PTTG1IP cleavage was located proximal to the fusion site (i.e. at the C-terminus). To take advantage of the natural cleavage of PTTG1IP, we generated a mutant whereby the terminal 50 amino acids (i.e. amino acids 130 and 180) were duplicated at the C-terminus to further promote Cre release (PTTG1IP-dup130-180) (**Fig 5a**).

**Figure 5.**
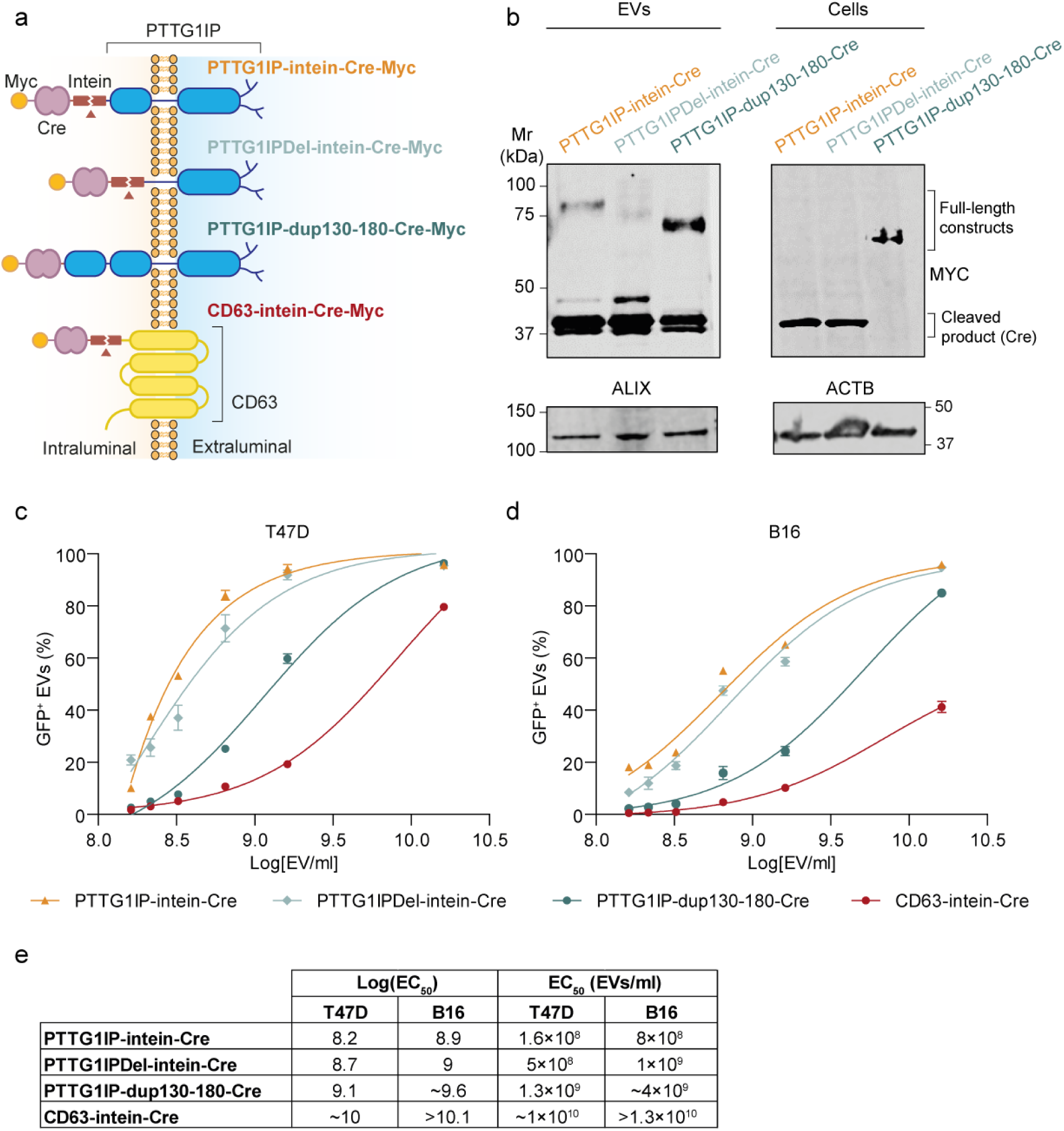
Engineered PTTG1IP variants exhibit superior EV-mediated delivery of Cre recombinase compared with equivalent CD63 constructs. HEK293T cells were transfected with various PTTG1IP and CD63 constructs and the capacity to transfer Cre protein in a dose response manner was tested in reporter cells. (**a**) Scheme of tested Cre fusion constructs. The intein is a self-cleaving sequence from Mycobacterium tuberculosis. PTTG1IPDel has a deletion from amino acid 130 to 164. PTTG1IP-dup130-180 has amino acids 130-180 duplicated on PTTG1IP C-terminus. (**b**) Western blot of Cre-fusion constructs in HEK293T cells and EVs. Both cell lysates and isolated EVs were blotted against MYC. Cell lysates and isolated EVs were blotted against ACTB and ALIX, respectively. (**c**) EV-transfer dose response in T47D cells. (**d**) EV-transfer dose response in B16 cells. Data are presented as mean ± SEM, *n* = 3 replicates. (**e**) LogEC50 and EC50 (EVs/ml) values of Cre fusion constructs.

To compare the efficiency of the various PTTG1IP variants (**Fig 5a,b)**, we performed an EV dose response experiment whereby B16 or T47D reporter cells were treated with EVs modified to express PTTG1IP-intein-Cre, PTTG1IP-dup130-180-Cre, PTTG1IP-Δ130-164-intein-Cre, or CD63-intein-Cre. A CD63-intein construct was included as a comparator scaffold as this tetraspanin has frequently been utilized for EV engineering purposes. PTTG1IP-intein-Cre and PTTG1IP1-Δ130-164-intein-Cre exhibited the highest potency followed by PTTG1IP-dup130-180- Cre (**Fig 5c,d**), while CD63-intein-Cre showed the lowest potency, especially in B16 cells (**Fig 5c,d**). Notably, PTTG1IP-intein-Cre was almost two logs more potent than CD63-intein-Cre in T47D cells, and at least one log more potent in B16 cells (**Fig 5e**). The higher potency of PTTG1IP-based scaffold may be at least partially explained by the higher EV-loading capacity of PTTG1IP as compared to CD63 (**Supplementary Fig 6**). The recombination efficiency of PTTG1IP-dup130-180-Cre EVs was double that of PTTG1IP-Cre at the same EV concentration (∼99% vs ∼40% in T47D cells), suggesting that Cre release was improved upon using the PTTG1IP-dup130-180 construct (**Fig 4e**, **Fig 5c,d**). This unexpected property may be beneficial as it could be exploited as a cargo release mechanism without the need for bacteria-derived cleaving sequences. Moreover, cleavage of PTTG1IP-dup130-180-Cre only occurred in the EVs. In contrast, for the PTTG1IP-intein-Cre construct, the cleaved Cre was readily detected in the donor cells (**Fig 5b**).

We next assessed if the cleavage sequence contained within PTTG1IP amino acids 130-180 could be utilised in other protein contexts. To this end, we compared the cleavage activity of the intein with the activity of a tandem-triplication of the EV-cleavage sequence derived from PTTG1IP, both of which were incorporated between CD63 and the Cre recombinase cargo (CD63-intein-Cre vs CD63-3xPTTG1IP130-180-Cre, **Supplementary Fig 7**). Cre cleavage in the EVs was observed with both PTTG1IP-dup130-180 and intein sequences, although the levels of cleavage were higher for the intein (**Supplementary Fig 7**). These data indicate that the PTTG1IP-derived intrinsic-cleavage sequence located at amino acids 130-180 can be used to promote cargo release in a distinct EV loading protein context.

### Assessment of Cre recombinase delivery to mice tumour xenografts

Building on positive results in cell cultures, we assessed EV-mediated Cre delivery *in vivo* using mice harbouring B16F10 Cre reporter cell tumour xenografts (**Fig 6a**). EVs were isolated from HEK293T cells transfected with PTTG1IP-intein-Cre + VSVG, Cre + VSVG, or VSVG alone and administered by intratumoral injection at a dose of 1×10^10^ EVs/mouse. Cre activity was subsequently analysed by PCR (using primers which span the reporter cassette) and immunohistochemistry at 4 days after EV treatment. Successful Cre recombination was detected in tumours harvested from the three PTTG1IP-intein-Cre treated mice with recombination efficiencies ranging from 46-90%, while no recombination was detected with Cre treated mice or controls (VSVG alone EVs or PBS) (**Fig 6b**). Similarly, GFP positive cells were detected in tumours from mice injected with PTTG1IP-intein-Cre, indicative of successful recombination, but not in tumours treated with passively loaded Cre (**Fig 6c**). In summary, these data suggest that PTTG1IP can be used as a scaffold for EV-mediated protein delivery *in vivo*.

**Figure 6.**
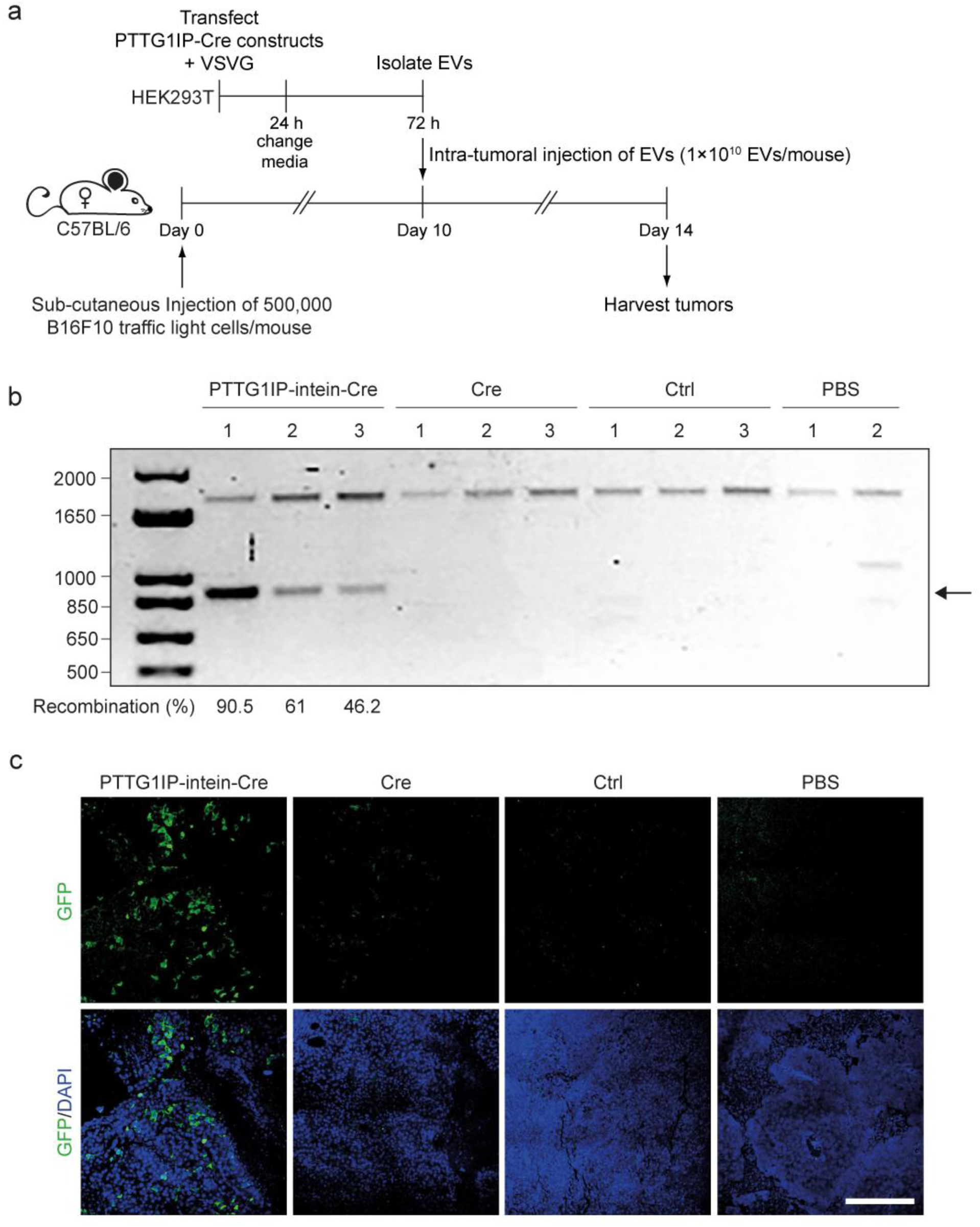
*In vivo* PTTG1IP-EV-mediated delivery of Cre recombinase. (**a**) Timeline of EV-transfer experiment to Cre reporter mice. 24h after donor cell transfection with Cre constructs and VSVG, media is replaced with fresh OptiMEM and EVs are isolated 48h later by SEC. 10 days before EV transfer, mice were injected with 500,000 B16F10 Cre reporter cells/mouse. On day 10, isolated EVs were injected to recipient mice by intratumoral injection (1×10^10^ EVs/mice). 4 days later, tumours are harvested and Cre activity is analysed. (**b**) Cre recombination activity in mice tumours treated with PTTG1IP-intein-Cre + VSVG, Cre + VSVG or VSVG only (control EVs). Cre recombination site was amplified by PCR and the products resolved by agarose gel electrophoresis. Cre recombination is indicated by a shorter 900 bp product (arrow) while non-recombined site is observed as a 1,700 bp product. Recombination percentages are indicated for recombined samples. (**c**) GFP expression in mice tumours treated with PTTG1IP-intein-Cre + VSVG EVs, Cre + VSVG EVs or PBS examined by ICH. *n* = 3 mice. Scale bar represents 150 µm.

### Assessment of Cas9 and Cas9/sgRNA complex delivery to reporter cells

Having shown that PTTG1IP can be used for loading and delivering cargo protein via EVs, we next assessed the ability of PTTG1IP to deliver more complex macromolecules, such as the CRISPR-Cas9 system. The CRISPR-Cas9 system represents a potent tool for gene editing and transcription regulation due to its high efficiency and specificity.^30^ However, this potential is hampered by the lack of efficient delivery methods.^31^ Thus, we explored the use of PTTG1IP to deliver Cas9 and Cas9/sgRNA complexes to recipient cells. For this purpose, we used HEK293T stoplight reporter cells.^17^ These cells express GFP upon Cas9 cleavage and indel formation (in 2 out of 3 of the editing events) and express RFP constitutively (**Fig 7a, Supplementary Fig 8**).

**Figure 7.**
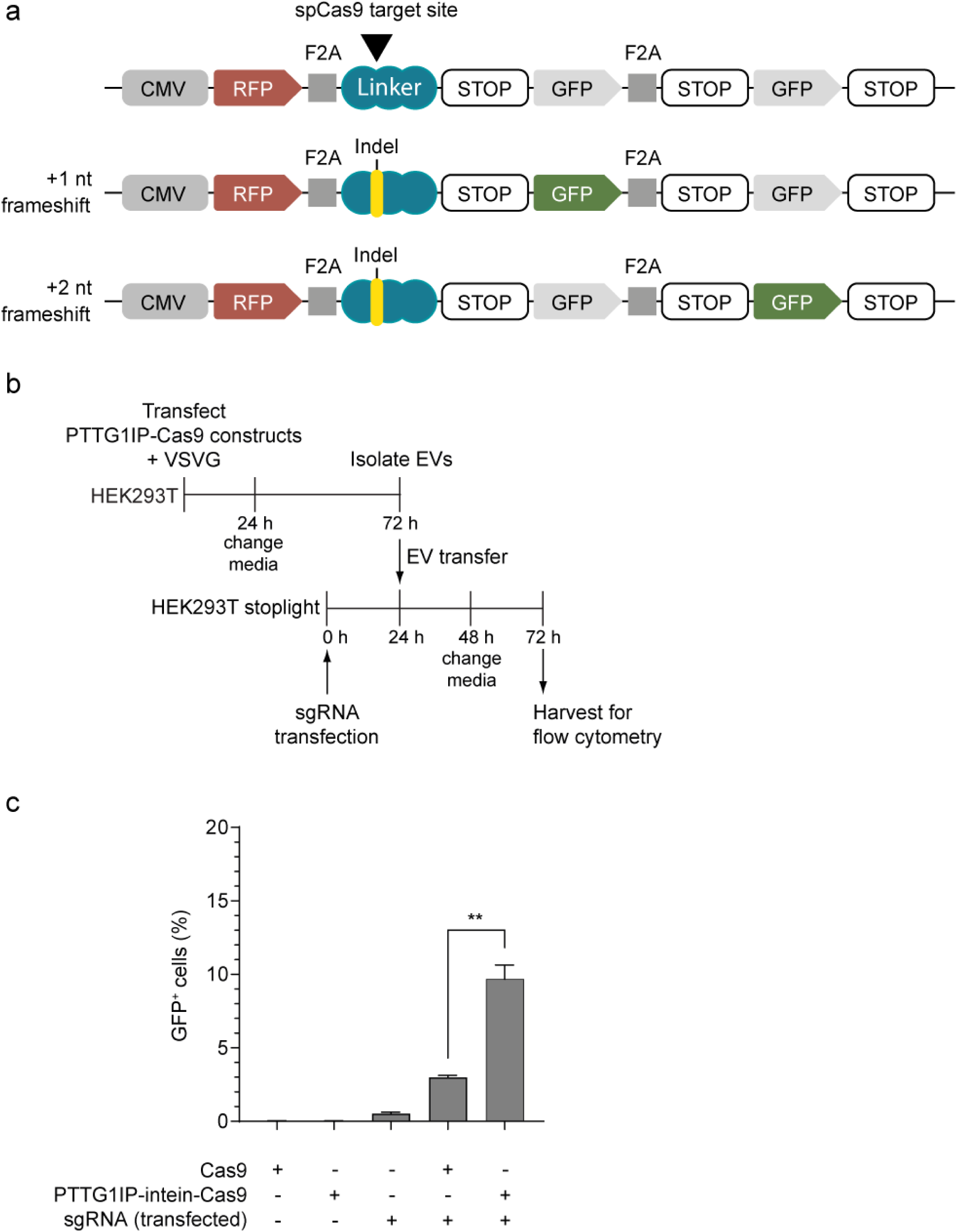
PTTG1IP-EV-mediated delivery of Cas9 protein. (**a**) Scheme of the Cas9 stoplight reporter construct. mCherry is constitutively expressed under a CMV promoter, followed by an F2A self-cleaving peptide sequence, a Cas9-binding linker region and a stop codon. Two GFP open reading frames that are frameshifted by one or two nucleotides are placed after the stop codon, separated by a F2A sequence. Upon Cas9 cleavage and indel formation, the translation reading frame will be shifted and GFP will be expressed together with mCherry (in 2/3 of the editing events). (**b**) Timeline of EV-transfer experiment. 24h after donor cell transfection with Cas9 constructs and VSVG, media is replaced with fresh Opti-MEM and EVs are isolated 48h later by SEC. Isolated EVs are then transferred to recipient cells (that were transfected with sgRNA 24h before) in 96-well plates (3×10^9^ EVs/ml). 24h later media is replaced with DMEM 10% FBS and after 24h cells are analyzed for GFP expression by flow cytometry. (**c**) GFP positive cells quantified by flow cytometry 48h after EV-transfer to sgRNA-transfected reporter cells. Statistical significance was tested by one-way ANOVA and Bonferroni *post hoc* test. Data are presented as mean + SEM, *n* = 3 replicates, \*\**P* < 0.01.

To assess Cas9 delivery, HEK293T cells were transfected with plasmids encoding Cas9 or PTTG1IP-intein-Cas9 together with VSVG expressing plasmid and EVs were isolated 72h later (**Fig 7b**). The recipient cells were pre-treated with sgRNA before EV transfer (so this experiment only assessed the delivery of Cas9 protein). 48h after EV treatment, Cas9 activity was assessed by flow cytometry to determine the percentage of GFP positive cells (**Fig 7b**). Activation after transfer of PTTG1IP-intein-Cas9 EVs occurred in ∼10% of the reporter cells, which was more than a 3-fold improvement as compared to unmodified Cas9 (**Fig 7c**). It should be noted that reporter cell activation after lipofection with Cas9 and sgRNA expressing plasmids was only ∼20% (**Supplementary Fig 8**). Moreover, Cas9 delivery was found to be twice as efficient with the PTTG1IP-intein-Cas9 scaffold than for CD63-intein-Cas9 (**Supplementary Fig 9**).

Having demonstrated the capacity to deliver Cas9 proteins, we were motivated to determine if PTTG1IP could be used to deliver Cas9/sgRNA RNP complexes. To assess Cas9-sgRNA complex delivery, reporter cells were treated with EVs from HEK293T cells transfected with Cas9 or PTTG1IP-intein-Cas9, together with VSVG and sgRNA (**Fig 8a**). Reporter cell activation using PTTG1IP-intein-Cas9 EVs was ∼3%, which was 6 times more efficient than activation with unmodified Cas9 EVs (**Fig 8b**). These results suggest that PTTG1IP is a suitable scaffold for delivery of more complex macromolecular cargoes.

**Figure 8.**
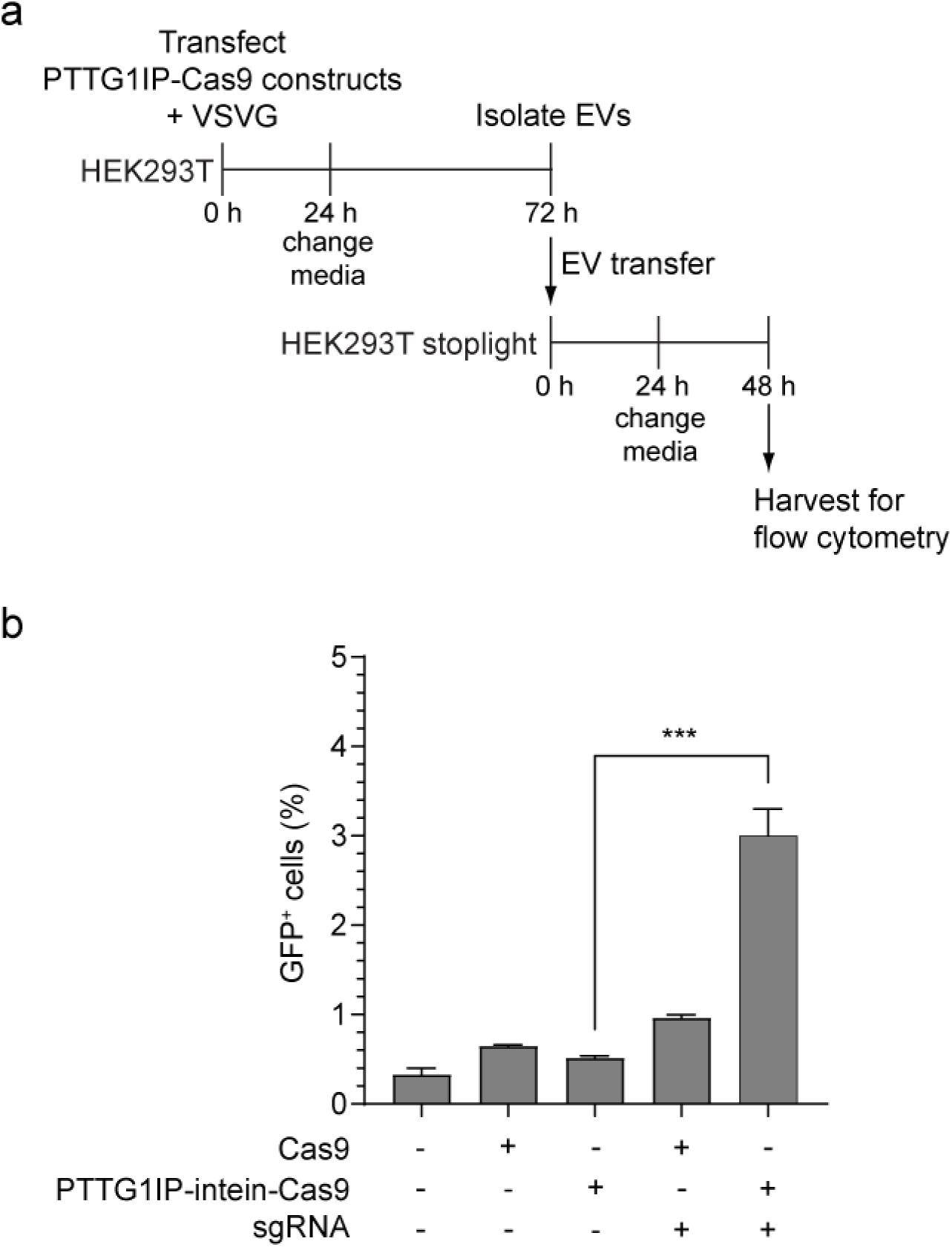
PTTG1IP-EV-mediated delivery of Cas9/sgRNA complexes. (**a**) Timeline of EV-transfer experiment to reporter cells. 24h after donor cell transfection with Cas9 constructs, sgRNA, and VSVG plasmids, media is replaced with fresh Opti-MEM and EVs are isolated 48h later by SEC. Isolated EVs are then transferred to recipient cells in 96-well plates (3×10^9^ EVs/ml). 24h later media is replaced with DMEM 10% FBS and after 24h cells are analyzed for GFP expression by flow cytometry. (**b**) GFP positive cells quantified by flow cytometry 48h after EV-transfer to reporter cells. Statistical significance was tested by one-way ANOVA and Bonferroni *post hoc* test. Data are presented as mean + SEM, *n* = 3 replicates, \*\*\**P* < 0.001.

### PTTG1IP engineering for surface display of targeting moieties

The topology of the PTTG1IP protein means that it is amenable to the addition of targeting moieties on its extracellular N-terminus via further protein engineering. To test this notion, a construct was developed in which the PTTG1IP scaffold was fused to a HER2 Single chain variable fragment (scFv) at the N-terminus,^32^ and a Nanoluciferase (Nluc) reporter at the C-terminus. EVs expressing this construct were harvested from HEK293T producer cells and incubated with the recipient SKBR3 human breast cancer cell line, which expresses HER2, and Nluc activity analyzed after 2 hours. Luciferase activity of recipient cells treated with HER2 scFv- PTTG1IP-Nluc EVs was higher than that of cells treated with control EVs without HER2 scFv (PTTG1IP-Nluc) at a dose of 2×10^6^ RLUs, although statistically significant (*P*=0.0429, one-tailed *t*-test) the effect was modest (**Supplementary** Figure 10). These data demonstrate the potential for further enhancement of PTTG1IP-based EV loading scaffolds.

## Discussion

A major challenge in developing EV-based therapies is the efficient loading of therapeutic cargo into the vesicles. Current methods for endogenous EV-protein loading often rely on tetraspanins,^33–35^ lipidic tags,^36–38^ or viral proteins (e.g. Gag),^39,40^ that despite showing a high loading capacity are not readily amenable for further engineering on the extracellular side (i.e. the addition of targeting or endosomal escape peptides). Lipidic tags and viral scaffolds target proteins to the lumen of the EVs, and engineering tetraspanins to display peptides or proteins on their extracellular loops while maintaining their EV-sorting capacity is non-trivial.^41–44^ To accomplish EV-loading and targeting, several strategies based on multi-modular EV-engineering have been developed,^45,46^ whereby distinct scaffold proteins are used to load the therapeutic cargo, and to display the targeting moiety. However, due to EV heterogeneity, there is a risk that these different modules (i.e. therapeutic cargo, targeting moiety, and endosomal escape moiety) are present in mutually exclusive EV subpopulations, thus diminishing the therapeutic potential of these EVs. There is therefore a need for more suitable EV scaffold proteins that allow for highly efficient intracellular EV-loading, and that are amenable to EV surface modification. Here, we have developed a bioinformatics analysis (IPFA) for identifying putative EV loading scaffolds based on PTMs (and other protein features), where N-glycosylated PTTG1IP was selected as the main candidate scaffold for EV cargo loading and functional delivery. We show that PTTG1IP EV-loading is dependent on its N-glycosylation status, and we demonstrate that PTTG1IP is able to efficiently deliver Cre protein *in vivo*, and Cas9/sgRNA complexes in cell culture. Furthermore, we present various PTTG1IP variants with improved properties with respect to their therapeutic application.

### Analysis of putative EV-sorting features

Candidate features identified with the IPFA analysis (PTAP motif, SPYR domain, MIT domain and transmembrane protein N-glycosylation) were assessed for EV-loading by fusing them to GFP.

To assess the role of N-glycosylation in EV-loading, we utilized PTTG1IP, the smallest protein with annotated N-glycosylation PTMs in our dataset. The PTTG1IP EV-scaffold showed the highest EV-loading capacity of any of the constructs tested (∼80% GFP^+^ vesicles) (**Supplementary Fig 5**).

PTTG1IP was shown to harbour two N-glycosylation sites at N54 (high mannose or hybrid N-glycosylation) and N45 (complex N-glycosylation), and its sorting to EVs was highly dependent on its N-glycosylation status, with the double glycosylation mutant showing greatly reduced EV-loading (∼27% GFP^+^ vesicles) than the single mutants (∼45% GFP^+^ vesicles) (**Fig 2**). Importantly, the cell expression levels of N-glycosylation mutants were similar (**Fig 2b,c**), suggesting that protein stability was not altered by loss of the N-glycosylation sites. These results demonstrate the robustness of IPFA analysis for identifying features that contribute to EV loading and identified PTTG1IP as a novel lead candidate scaffold.

### EV-loading of PTTG1IP variants

PTTG1IP is a transmembrane protein that is ubiquitously expressed in normal human tissues.^47^ PTTG1IP is conserved across a wide variety of animal species and does not share significant homology with other human proteins.^48^ Initial studies on PTTG1IP function suggested a role in trafficking pituitary tumor-transforming gene 1 (PTTG) to the nucleus, where PTTG induces tumorigenesis via various mechanisms including transcription activation and p53 destabilization.^47,49^ However, subsequent studies have shown that PTTG1IP is primarily located on intracellular vesicles structures and not in the nucleus.^48,50^ Both PTTG and PTTG1IP are upregulated in several types of cancer,^51,52^ and it has been reported that PTTG1IP has an independent transforming ability by inducing p53 degradation.^53^ Conversely, PTTG1IP downregulation has been reported in lung cancer tissues^54^ and has been associated with a higher risk of breast cancer death.^55^ In addition, PTTG1IP overexpression was shown to inhibit proliferation in lung cancer cells.^54^

Although there is conflicting evidence relating to the pro-oncogenic potential of PTTG1IP, we were interested to determine if the putative p53 interacting domain could be deleted without affecting EV loading potential. To this end, we deleted the coiled coil region at the PTTG1IP C-terminus (amino acids 130 – 164). This region contains the NLS (nuclear localization signal) bipartite signal that is required for its interaction with PTTG^47^ as well as the potential p53 binding site.^49^ PTTG1IP 130-164 deletion retained EV-sorting capacity, suggesting that the deleted region is not required for EV-sorting (**Fig 3c,d**). Furthermore, this PTTG1IP deletion variant was almost as potent as the WT PTTG1IP scaffold when assessed for Cre delivery (**Fig 5c,d**). In this manner, potential interactions between PTTG1IP and p53 could be avoided through careful protein engineering.

Mathieu *et al*., reported that a CD63 endocytosis signal (YXXΦ) mutant was more abundantly released in the EVs than WT CD63.^24^ This increase was due to a re-direction of CD63 from being released in exosomes via the MVB to being released in microvesicles originating from the plasma membrane. Thus, we mutated the two endocytosis signals on PTTG1IP (2YA mutant) to determine if EV-loading could be improved. In line with the previous study, PTTG1IP 2YA mutant EVs were able to load twice as much GFP per vesicle relative to WT PTTG1IP EVs (**Fig 3c,d**). It is possible that the mechanism by which CD63 loading is increased also applies to PTTG1IP, although this hypothesis was not further explored. PTTG1IP could be further engineered to optimize its utility for EV delivery. For example, future studies could test the effect of introducing additional N-glycosylation moieties on PTTG1IP to increase EV-loading efficiency. In addition, PTTG1IP exhibits a remarkable tolerance to relatively large internal deletions. Further development could enhance its ‘scaffold-like’ properties (e.g. engineering the extracellular domain to display targeting or cell signalling moieties) while diminishing the native functions of PTTG1IP (e.g. intracellular domain modification).

### EV-mediated delivery of PTTG1IP-Cre

To determine if PTTG1IP scaffolds could be used to functionally deliver EV-protein cargo, we assessed Cre packaging and delivery to reporter cells. To promote Cre release in the EVs, we used two different self-cleaving sequences that have been reported to cleave at low pH conditions similar to those in the endosomal compartment.^26,28^ In addition, we co-expressed VSVG in the EV-producing cells to promote EV endosomal escape in the recipient cells. We showed that PTTG1IP promoted EV-packaging of Cre and that cleavage by the intein sequence was much more efficient than with the SC sequence (**Fig. 4b**). Importantly, PTTG1IP was able to deliver Cre to a variety of reporter cell lines with high efficiency when the intein was used as the release sequence, and VSVG was co-expressed to promote endosomal escape (∼85-99%) (**Fig 4e,f**, and **Fig 5d,e**). The higher potency of PTTG1IP as compared to CD63 scaffold may be partially explained by the higher (∼30%) EV-loading capacity of the former (**Supplementary Fig 6**). However, the magnitude of this increase does not seem to be enough to fully explain this phenomenon. It is possible that there are a higher proportion of PTTG1IP^+^ VSVG^+^ double positive EVs than CD63^+^ VSVG^+^ EVs. Alternatively, PTTG1IP may somehow enhance EV uptake by recipient cells.

Notably, some degree of Cre cleavage was detected in the EVs from PTTG1IP-Cre fusion protein when no exogenous self-cleavage domain was present (**Fig 4b**). Treatment with PTTG1IP-Cre also resulted in reporter cell activation, although not as efficiently as PTTG1IP-intein-Cre (**Fig 4e,f**). These data suggested that PTTG1IP exhibits an intrinsic cleavage potential although this is much less efficient than the intein sequence. Duplication of the potential cleaving sequence in PTTG1IP resulted in a two-fold Cre delivery improvement (**Fig 4e** and **Fig 5c**), suggesting that a cleavage site exists between amino acids 130-180 of PTTG1IP. Importantly, we also showed that the sequence could be exploited to induce EV-cleavage in different protein contexts such as CD63 (**Supplementary Fig 7**). We hypothesize that this sequence contains a putative enzymatic cleavage site, and that cleavage predominantly occurs in the EVs (**Fig 5b**). The PTTG1IP intrinsic cleavage sequence presents several advantages over existing cargo release systems for future construct engineering. Firstly, PTTG1IP cleavage sequence is human and thus most likely non-immunogenic, as opposed to the intein sequence, which is derived from *Mycobacterium tuberculosis*. A potential B-cell response towards intein-derived peptides may be avoided when EVs are generated externally by producer cells given that the sequence will be hidden inside the EVs. However, if the EVs are to be produced endogenously *in situ* as previously shown,^46,56^ the expression of a bacterial peptide would most likely trigger an immune response towards the producer cells and its subsequent elimination, hampering its therapeutic application. Secondly, PTTG1IP-Cre cleavage was only detected in the EVs while intein cleavage was detected in the EV producer cells. This feature may also be valuable to maximize cargo packaging, since cargo release from the PTTG1IP EV-scaffold before loading into the EVs would be minimal. Previous studies have relied on blue-light/small molecule-induced dimerization,^33,57^ or on the introduction of viral cleavable linkers derived from murine leukemia virus (MLV)^39^ or HIV-1^58^ to release protein cargo from the EV-loading scaffolds. However, these strategies present several issues. Conditional dimerization strategies are based on the production of EVs under blue light or the presence of a small molecule that promotes dimerization of the loading scaffold and the protein cargo. Hence, these strategies cannot be applied to *in situ* EV therapies. Moreover, small molecules used to induce dimerization will also be delivered to recipient cells. Viral cleavable sequences require the expression of viral proteases in the donor cells, which besides being immunogenic may be packaged into EVs and delivered to recipient cells. Thus, once optimized, the PTTG1IP cleavage sequence would constitute a safer and more convenient alternative for EV-based therapies.

*In vivo* Cre delivery was demonstrated using EVs expressing PTTG1IP-intein-Cre and VSVG in mice harboring B16 reporter cell tumor xenografts (**Fig 6**). Recombination efficiency ranged from 40% to 90%, conversely, no activation was detected in mice treated with Cre and VSVG EVs, suggesting that active loading is required for Cre delivery.

### EV-mediated delivery of Cas9 to reporter cells

Delivery of Cas9 using PTTG1IP as scaffold and the intein for cargo release enabled the activation of ∼10% of Cas9 reporter cells (**Fig 7**), while conventional transfection of Cas9 and sgRNA resulted in ∼20% activation (**Supplementary Fig 8**). In line with Cre delivery results, PTTG1IP was two times more efficient in delivering Cas9 as compared to CD63 scaffold (**Supplementary Fig 9**). PTTG1IP-intein strategy was also able to successfully deliver Cas9/sgRNA ribonucleoprotein complex to reporter cells, although the delivery efficiency was reduced as compared to Cas9 protein transfer alone. Both Cas9 and Cas9/sgRNA delivery are complex processes involving two different components and thus a low delivery efficiency was expected. Cas9 delivery requires binding of Cas9 to the sgRNA in the nucleus after the Cas9 is delivered into the cytosol, while for Cas9/sgRNA delivery the sgRNA must be exported from the nucleus to bind Cas9 in the cytoplasm before it is sorted to the EVs. In addition, the sensitivity of the reporter cells was low, as observed by the activation level after Cas9 and sgRNA lipofection (**Supplementary Fig 8**).

Other studies have attempted to load and deliver Cas9/sgRNA complex via EVs with variable degrees of success. Initial studies relied on centrifugation of EVs together with recipient cells (spinoculation) in order to deliver Cas9/sgRNA cargo.^59,60^ However, despite the uptake efficiency of this method being high, it can only be used for *in vitro* delivery. Cas9/sgRNA sorting to EVs has also been achieved by simple overexpression of dCas9-VPR protein and sgRNA.^61,62^

Nevertheless, in this and other studies, functional delivery of Cas9 was not detected in EVs unless Cas9 was tagged with an EV-targeting protein.^5,63,64^ Several recent studies have successfully delivered Cas9 ribonucleoprotein using Gag viral proteins,^39,64^ ARRDC1,^65^ or CD63^5,66^ as EV- scaffold proteins. In most of these studies, a release system was required to accomplish functional delivery of Cas9. These were based on aptamer-aptamer binding protein interaction, GFP-GFP nanobody interaction, small molecule induced dimerization or the introduction of a viral protease restriction site. This was not the case for the strategy based on ARMMs (arrestin domain containing protein1-mediated microvesicles), in which ARMMs were capable of packaging and delivering Cas9/sgRNA complex *in vitro* with moderate efficiency when Cas9 was fused to ARRDC1-interacting WW-domains.^65^ Another study used a myristoylation tag on Cas9 to successfully deliver Cas9/sgRNA complex to recipient cells *in vitro*.^14^ However, chloroquine and polybrene transfection reagent were added during the EV transfer, limiting the therapeutic use of this strategy.^14^ As for the requirement of an endosomal escape protein, some studies have also included VSVG^2,29,62,67–69^ or BaEVRLess^39^ (baboon endogenous virus envelope glycoprotein) at the EV surface to enhance uptake and cytoplasmic delivery. Direct comparison of our study with other EV-based Cas9/sgRNA delivery studies can be challenging given that the EV concentration and gene editing efficiency have frequently not been reported in many of the studies. Moreover, efficiency of EV delivery may depend on the EV isolation method, donor-recipient cell line combinations, and reporter system.

The PTTG1IP-based platform offers several advantages over existing protein scaffolds for EV- delivery. We have demonstrated PTTG1IP is able to efficiently deliver protein cargo in cell culture and *in vivo.* EV-scaffolds based on tetraspanins, lipidic tags, VSVG, Gag, or ARRDC1 are not readily amenable to further engineering to display targeting or endosomal escape peptides on the EV surface. Hence a second scaffold that co-localizes with the first should be used for extracellular cargo loading, which would be difficult to accomplish due to vesicle heterogeneity.

Unlike VSVG and Gag viral scaffolds, PTTG1IP would presumably be less immunogenic given that it is an endogenous protein. In addition, PTTG1IP is able to package protein cargoes with high efficiency, in contrast with alternative single spanning proteins (i.e. LAMP2B, PDGFR, and IGFS8) that exhibit lower EV-loading than CD63.^41,70^ Recently, Dooley *et al*., identified single spanning protein PTFGRN as a highly EV-enriched protein that enables efficient protein surface display.^70^ Although PTFGRN could potentially be used for delivery of intraluminal cargoes, this has yet to be demonstrated. In contrast to scaffolds that rely on EV-surface association (i.e. CP05 peptide,^45^ C1C2 domain derived from lactadherin^71^, GPI-anchor,^37^ and G58 peptide derived from GAPDH),^71^ cargo molecules fused to the PTTG1IP C-terminus are protected from degradation since cargo will not be exposed to the extracellular milieu.

The PTTG1IP based platform described here has been shown to be able to efficiently deliver macromolecular cargoes *in vitro* and *in vivo*. Nevertheless, there is scope for further optimization to fully harness its potential. Most importantly, our system requires VSVG for functional delivery, thus future studies should evaluate human-derived fusogenic proteins to substitute for VSVG. We have demonstrated that PTTG1IP can tolerate the addition of a targeting moiety (i.e. scFv targeting HER2) at its extracellular terminus. Substitution of this targeting module with other targeting ligands may enable more efficient cellular uptake and/or delivery to specific target cells or tissues. In addition, the suitability of PTTG1IP to display peptide moieties or protein cargo on the EV surface should also be assessed. Further engineering of the PTTG1IP extracellular domain has the potential to be used for EV surface display (e.g., receptor decoys, fusogenic peptides or signalling proteins), which would allow for both extra- and intraluminal EV modification. Moreover, the PTTG1IP cleavage sequence should be further characterized and optimized in order to achieve cleavage levels similar to, or better than, those of the intein.

## Conclusion

In summary, we describe the development of a highly versatile platform for EV-mediated delivery of protein and protein/sgRNA complexes based on an N-glycosylated PTTG1IP scaffold. We also demonstrate the utility of IPFA analysis for the identification of EV-enriched features, and we show that protein PTMs can be exploited to enhance functional cargo loading into EVs. Thus, our study expands the range of tools available for engineering EVs into efficient vehicles for targeted delivery of therapeutics.

## Author Contributions

This work was conceived by CMP, IM, and TCR. CMP, XL, and DG performed experiments and collected data. All authors were involved in data analysis. IM, SEA, MJAW, and TCR supervised the work. CMP and TCR wrote the first draft of the manuscript. All authors contributed to the final draft of the manuscript.

## Acknowledgements

This work was supported by an MRC Programme Grant awarded to MJAW (MRN0248501) and John Fell Fund grant awarded to TCR. CM, IM, and MJAW are supported by Horizon 2020 (B- SMART). SEA is supported by the following grants: Horizon 2020 (EXPERT), the Swedish foundation of Strategic Research (FormulaEx, SM19-0007), ERC CoG (DELIVER), and the Swedish Research Council (4–258/2021).

