## Supplementary Information for "An extracellular vesicle delivery platform based on the PTTG1IP protein"

| Application | Antibody | Host | ID | Manufacturer | Dilution |
|---|---|---|---|---|---|
| Primary | Anti-GFP | Rabbit | ab290/ 338001 | Abcam/Biolegend | 1:1,000 |
| Primary | Anti-cMyc | Mouse | 626801 | Biolegend | 1:1,000 |
| Primary | Anti-CD63 | Mouse | Ab1289 | Abcam | 1:1,000 |
| Primary | Anti-ACTB | Mouse | MA5-15739 | ThermoFisher | 1:2,000 |
| Primary | Anti-ALIX | Rabbit | ab186429 | Abcam | 1:2,000 |
| Secondary | Anti-Rabbit | Goat | 926-32211 | LI-COR | 1:5,000 |
| Secondary | Anti-Mouse | Goat | 926-32210 | LI-COR | 1:5,000 |

**Supplementary Table 1**

**Details of antibodies used in this study.**

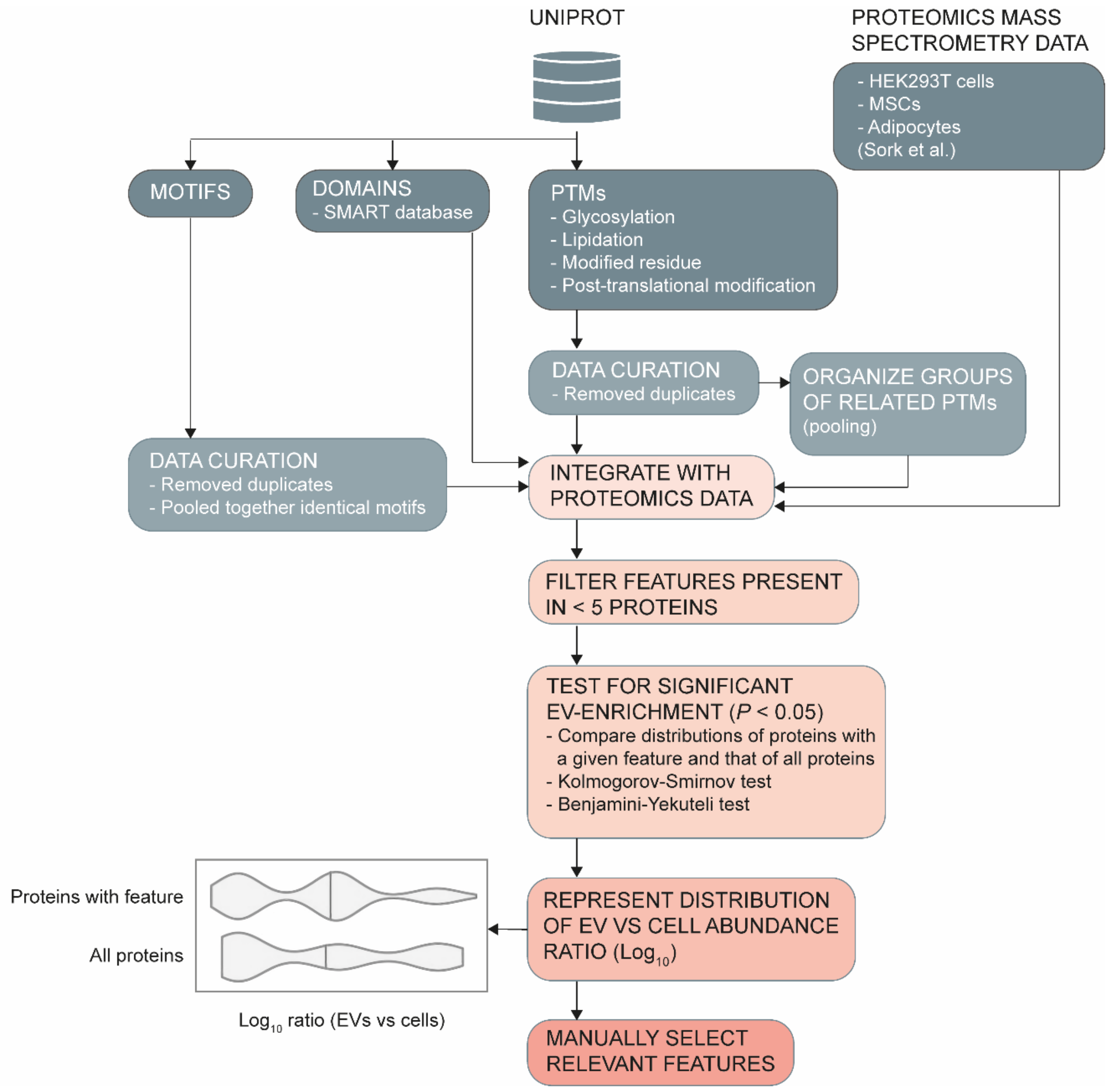
UNIPROT
PROTEOMICS MASS SPECTROMETRY DATA
- HEK293T cells
- MSCs
- Adipocytes
(Sork et al.)
MOTIFS
DOMAINS
- SMART database
PTMs
- Glycosylation
- Lipidation
- Modified residue
- Post-translational modification
DATA CURATION
- Removed duplicates
ORGANIZE GROUPS OF RELATED PTMs
(pooling)
DATA CURATION
- Removed duplicates
- Pooled together identical motifs
INTEGRATE WITH PROTEOMICS DATA
FILTER FEATURES PRESENT IN < 5 PROTEINS
TEST FOR SIGNIFICANT EV-ENRICHMENT (P < 0.05)
- Compare distributions of proteins with a given feature and that of all proteins
- Kolmogorov-Smirnov test
- Benjamini-Yekuteli test
Proteins with feature
All proteins
Log10 ratio (EVs vs cells)
REPRESENT DISTRIBUTION OF EV VS CELL ABUNDANCE RATIO (Log10)
MANUALLY SELECT RELEVANT FEATURES

**Supplementary Figure 1**

**Bioinformatics analysis scheme.**

Protein motif, domain (as annotated on SMART database), and post-translational modification (PTM) data were extracted from the UniProt database. Motif and PTM data were then manually curated to remove duplicates and pool together identical features. In the case of PTMs, additional groups of related PTMs were pooled and included in the analysis as separate categories. Protein feature data were integrated with mass spectrometry proteomics data325 (from HEK293T cells, MSCs, and adipocytes) and only features that appear on at least 5 proteins were included in the analysis because of the difficulty to detect statistically significant changes of feature annotations when data was scant. Distributions of proteins with a given feature and that of all proteins were then compared by Kolmogorov-Smirnov test followed by Benjamini-Yekuteli *post hoc* test to identify features that were significantly EV-enriched ($P < 0.05$). The distribution of EV vs cell abundance ratio ($\log_{10}$) of proteins with significantly EV-enriched features was then represented and relevant features were manually selected.

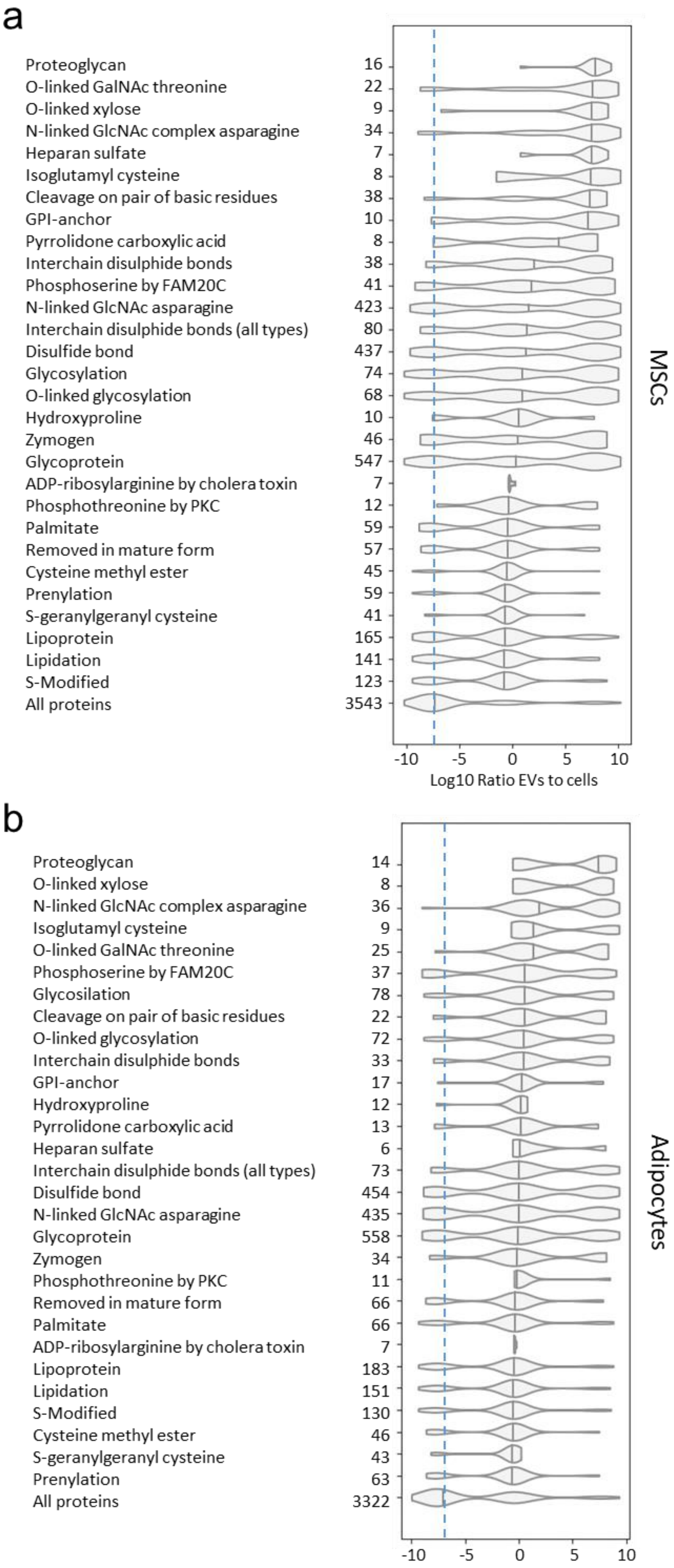
a
Proteoglycan 16
O-linked GalNAc threonine 22
O-linked xylose 9
N-linked GlcNAc complex asparagine 34
Heparan sulfate 7
Isoglutamyl cysteine 8
Cleavage on pair of basic residues 38
GPI-anchor 10
Pyrrolidone carboxylic acid 8
Interchain disulphide bonds 38
Phosphoserine by FAM20C 41
N-linked GlcNAc asparagine 423
Interchain disulphide bonds (all types) 80
Disulfide bond 437
Glycosylation 74
O-linked glycosylation 68
Hydroxyproline 10
Zymogen 46
Glycoprotein 547
ADP-ribosylarginine by cholera toxin 7
Phosphothreonine by PKC 12
Palmitate 59
Removed in mature form 57
Cysteine methyl ester 45
Prenylation 59
S-geranylgeranyl cysteine 41
Lipoprotein 165
Lipidation 141
S-Modified 123
All proteins 3543
MSCs
-10 -5 0 5 10
Log10 Ratio EVs to cells
b
Proteoglycan 14
O-linked xylose 8
N-linked GlcNAc complex asparagine 36
Isoglutamyl cysteine 9
O-linked GalNAc threonine 25
Phosphoserine by FAM20C 37
Glycosilation 78
Cleavage on pair of basic residues 22
O-linked glycosylation 72
Interchain disulphide bonds 33
GPI-anchor 17
Hydroxyproline 12
Pyrrolidone carboxylic acid 13
Heparan sulfate 6
Interchain disulphide bonds (all types) 73
Disulfide bond 454
N-linked GlcNAc asparagine 435
Glycoprotein 558
Zymogen 34
Phosphothreonine by PKC 11
Removed in mature form 66
Palmitate 66
ADP-ribosylarginine by cholera toxin 7
Lipoprotein 183
Lipidation 151
S-Modified 130
Cysteine methyl ester 46
S-geranylgeranyl cysteine 43
Prenylation 63
All proteins 3322
Adipocytes
-10 -5 0 5 10

**Supplementary Figure 2**

***In silico*-identified EV-enriched PTM annotations in MSCs and adipocytes.**

Violin plots representing the EV to cell abundance ratios ($log_{10}$) of proteins with statistically significant EV-enriched PTMs identified in (**a**) MSCs and (**b**) adipocytes. Only EV-enriched annotated PTMs common across all three cell lines are shown. Numbers on the y-axes depict the number of proteins with the given PTM in the dataset. Violin plot of the EV to cell abundance ratios ($log_{10}$) of all proteins in the dataset is included as a reference. Median value of all proteins in the dataset is represented as a dashed line. $^*P < 0.05$.

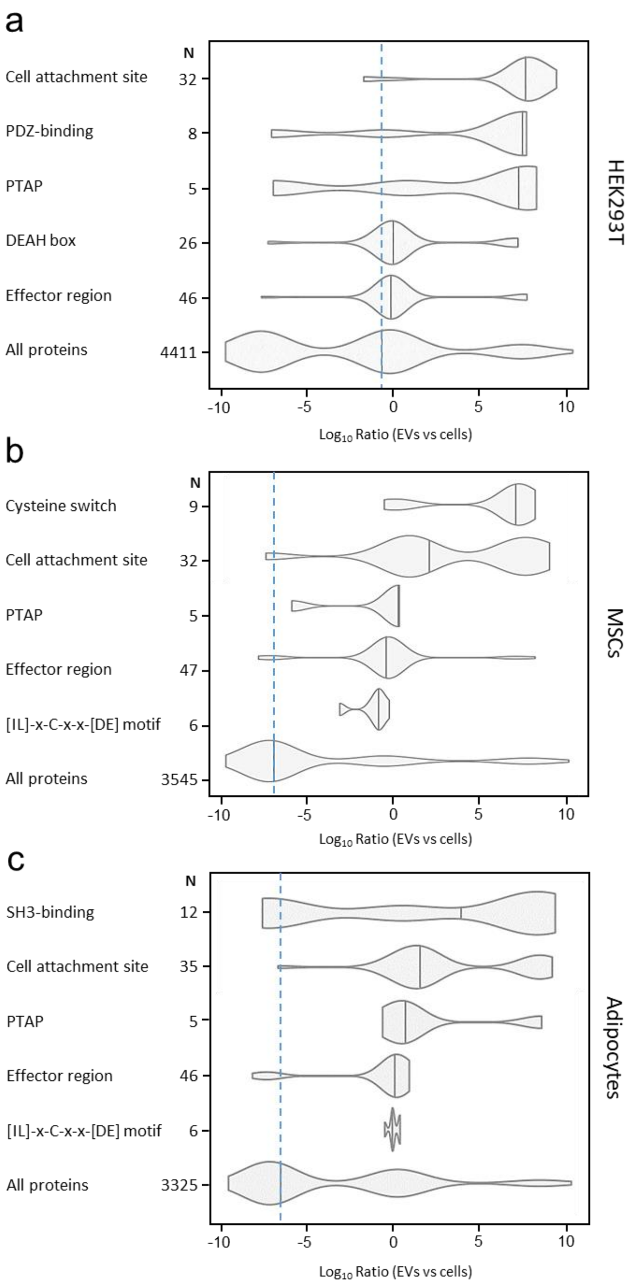
a
N
Cell attachment site 32
PDZ-binding 8
PTAP 5
DEAH box 26
Effector region 46
All proteins 4411
HEK293T
-10 -5 0 5 10
Log10 Ratio (EVs vs cells)
b
N
Cysteine switch 9
Cell attachment site 32
PTAP 5
Effector region 47
[IL]-x-C-x-x-[DE] motif 6
All proteins 3545
MSCs
-10 -5 0 5 10
Log10 Ratio (EVs vs cells)
c
N
SH3-binding 12
Cell attachment site 35
PTAP 5
Effector region 46
[IL]-x-C-x-x-[DE] motif 6
All proteins 3325
Adipocytes
-10 -5 0 5 10
Log10 Ratio (EVs vs cells)

**Supplementary Figure 3**

***In silico*-identified EV-enriched annotated motifs.**

Violin plots representing the EV to cell abundance ratios ($log_{10}$) of proteins with statistically significant EV-enriched annotated motifs identified in HEK293T cells (**a**), MSCs (**b**) and adipocytes (**c**). Numbers on the y-axes depict the number of proteins with the given motif in the dataset. Violin plot of the EV to cell abundance ratios ($log_{10}$) of all proteins in the dataset is included as a reference. Median value of all proteins in the dataset is represented as a dashed line. $^*P < 0.05$.

a

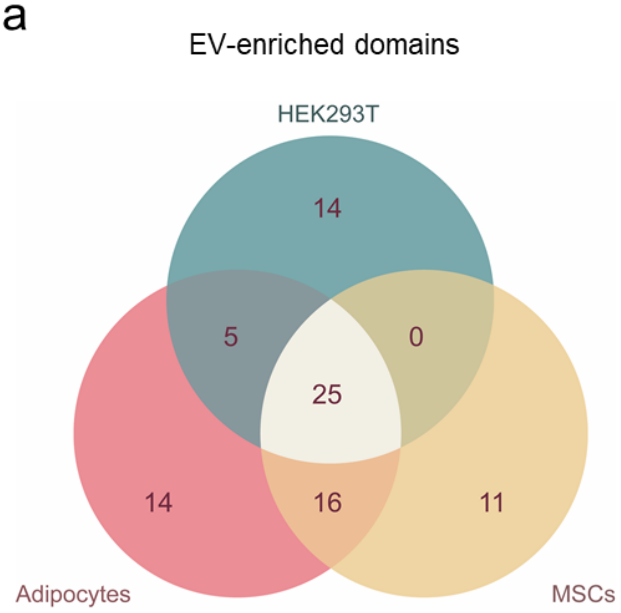
EV-enriched domains
HEK293T
14
5
0
25
14
16
11
Adipocytes
MSCs

b

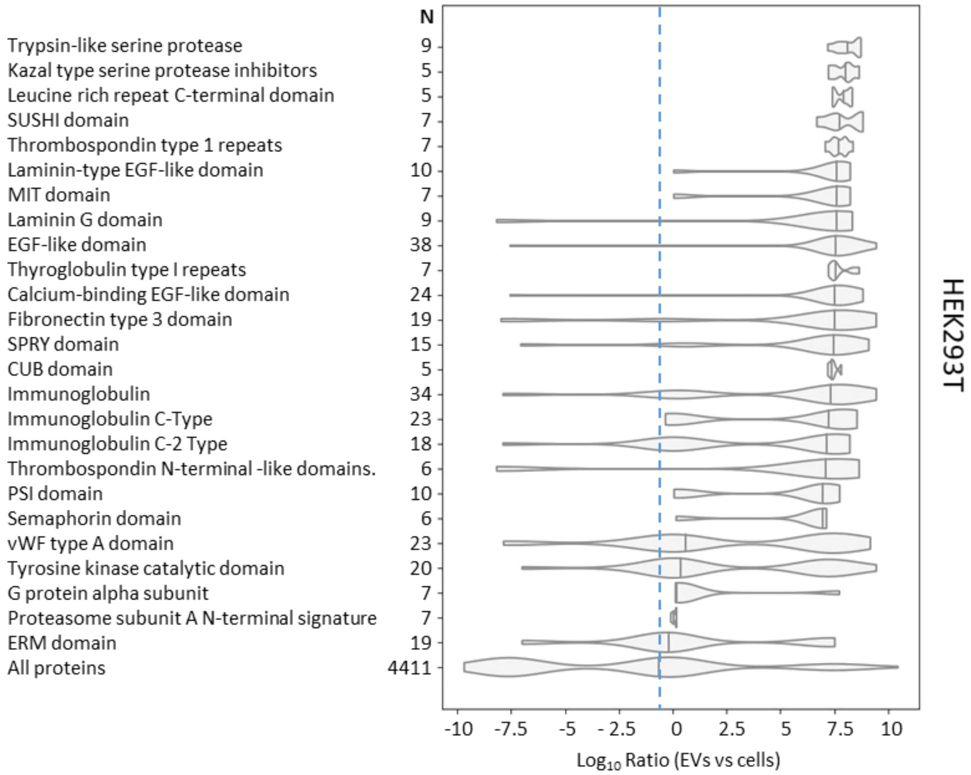
N
Trypsin-like serine protease 9
Kazal type serine protease inhibitors 5
Leucine rich repeat C-terminal domain 5
SUSHI domain 7
Thrombospondin type 1 repeats 7
Laminin-type EGF-like domain 10
MIT domain 7
Laminin G domain 9
EGF-like domain 38
Thyroglobulin type I repeats 7
Calcium-binding EGF-like domain 24
Fibronectin type 3 domain 19
SPRY domain 15
CUB domain 5
Immunoglobulin 34
Immunoglobulin C-Type 23
Immunoglobulin C-2 Type 18
Thrombospondin N-terminal -like domains. 6
PSI domain 10
Semaphorin domain 6
vWF type A domain 23
Tyrosine kinase catalytic domain 20
G protein alpha subunit 7
Proteasome subunit A N-terminal signature 7
ERM domain 19
All proteins 4411
-10 -7.5 -5 - 2.5 0 2.5 5 7.5 10
Log10 Ratio (EVs vs cells)
HEK293T

c

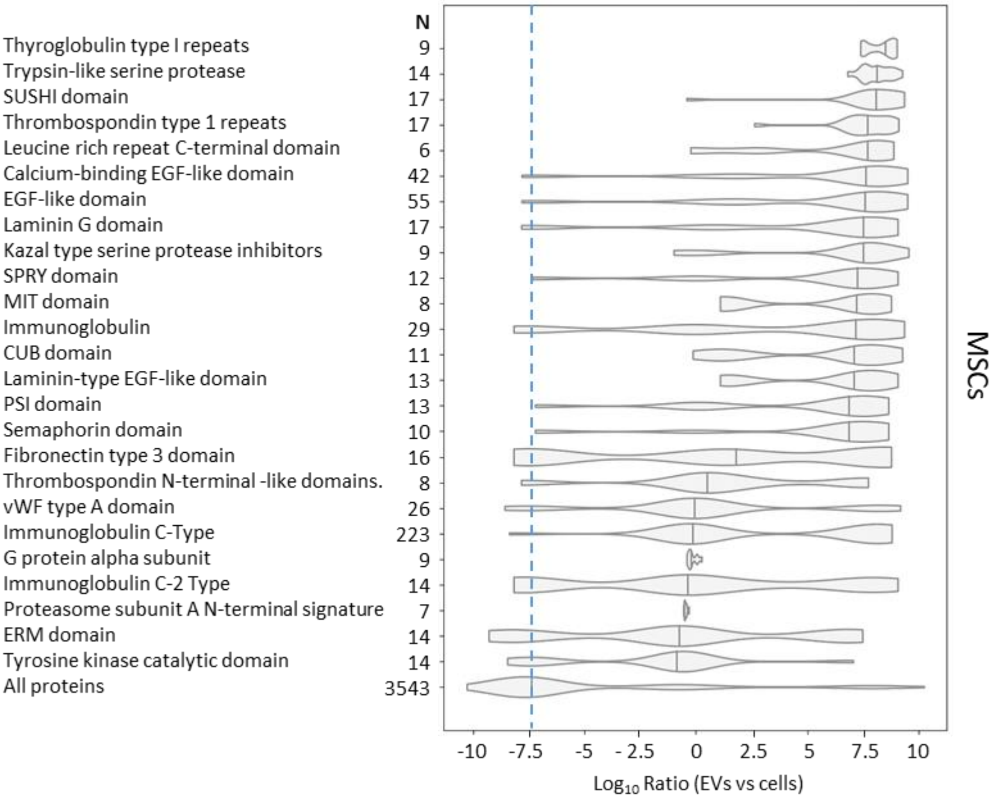
N
Thyroglobulin type I repeats 9
Trypsin-like serine protease 14
SUSHI domain 17
Thrombospondin type 1 repeats 17
Leucine rich repeat C-terminal domain 6
Calcium-binding EGF-like domain 42
EGF-like domain 55
Laminin G domain 17
Kazal type serine protease inhibitors 9
SPRY domain 12
MIT domain 8
Immunoglobulin 29
CUB domain 11
Laminin-type EGF-like domain 13
PSI domain 13
Semaphorin domain 10
Fibronectin type 3 domain 16
Thrombospondin N-terminal -like domains. 8
vWF type A domain 26
Immunoglobulin C-Type 223
G protein alpha subunit 9
Immunoglobulin C-2 Type 14
Proteasome subunit A N-terminal signature 7
ERM domain 14
Tyrosine kinase catalytic domain 14
All proteins 3543
MSCs
-10 -7.5 -5 - 2.5 0 2.5 5 7.5 10
Log10 Ratio (EVs vs cells)

d

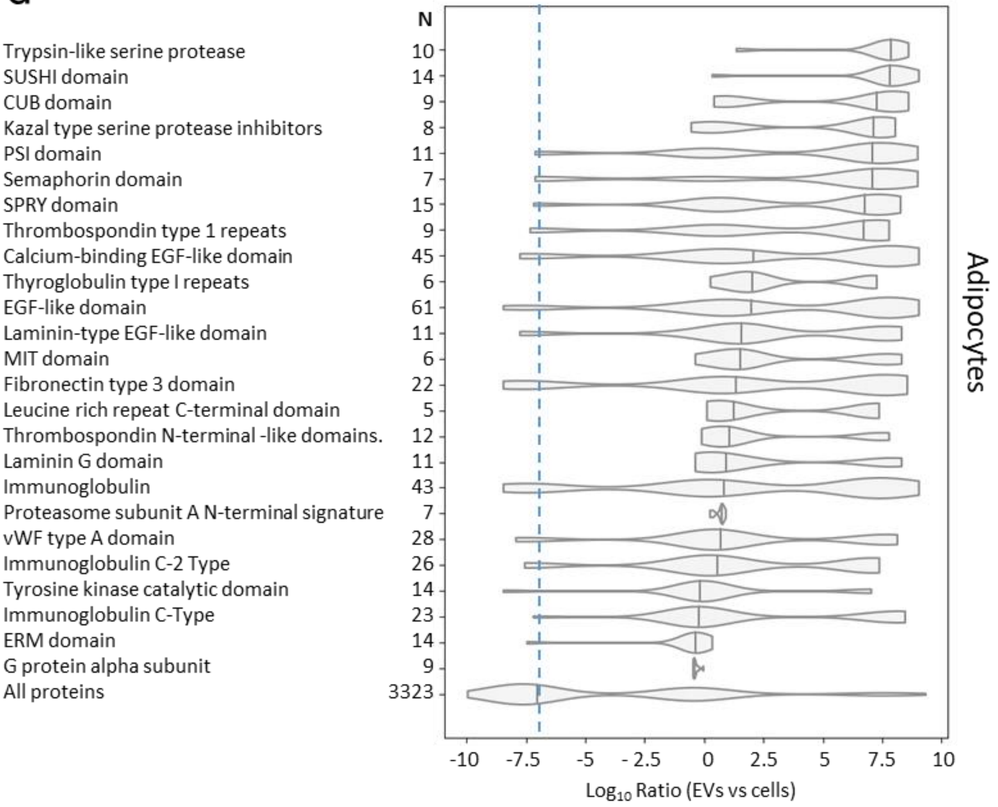
N
Trypsin-like serine protease 10
SUSHI domain 14
CUB domain 9
Kazal type serine protease inhibitors 8
PSI domain 11
Semaphorin domain 7
SPRY domain 15
Thrombospondin type 1 repeats 9
Calcium-binding EGF-like domain 45
Thyroglobulin type I repeats 6
EGF-like domain 61
Laminin-type EGF-like domain 11
MIT domain 6
Fibronectin type 3 domain 22
Leucine rich repeat C-terminal domain 5
Thrombospondin N-terminal -like domains. 12
Laminin G domain 11
Immunoglobulin 43
Proteasome subunit A N-terminal signature 7
vWF type A domain 28
Immunoglobulin C-2 Type 26
Tyrosine kinase catalytic domain 14
Immunoglobulin C-Type 23
ERM domain 14
G protein alpha subunit 9
All proteins 3323
Adipocytes
-10 -7.5 -5 - 2.5 0 2.5 5 7.5 10
Log10 Ratio (EVs vs cells)

**Supplementary Figure 4**

***In silico*-identified EV-enriched domain annotations.**

(**a**) Venn diagram of statistically significant EV-enriched annotated domains in HEK293T cells, MSCs and adipocytes. Numbers in the diagram refer to the number of domains identified for each group. Violin plots representing the EV to cell abundance ratios ($\log_{10}$) of proteins with statistically significant EV-enriched domains identified in HEK293T cells (**b**), MSCs (**c**) and adipocytes (d). Only EV-enriched annotated domains common across all three cell lines are shown. Numbers on the y-axes depict the number of proteins with the given domain in the dataset. Violin plot of the EV to cell abundance ratios ($\log_{10}$) of all proteins in the dataset is included as a reference. Median value of all proteins in the dataset is represented as a dashed line. *$P$ < 0.05.

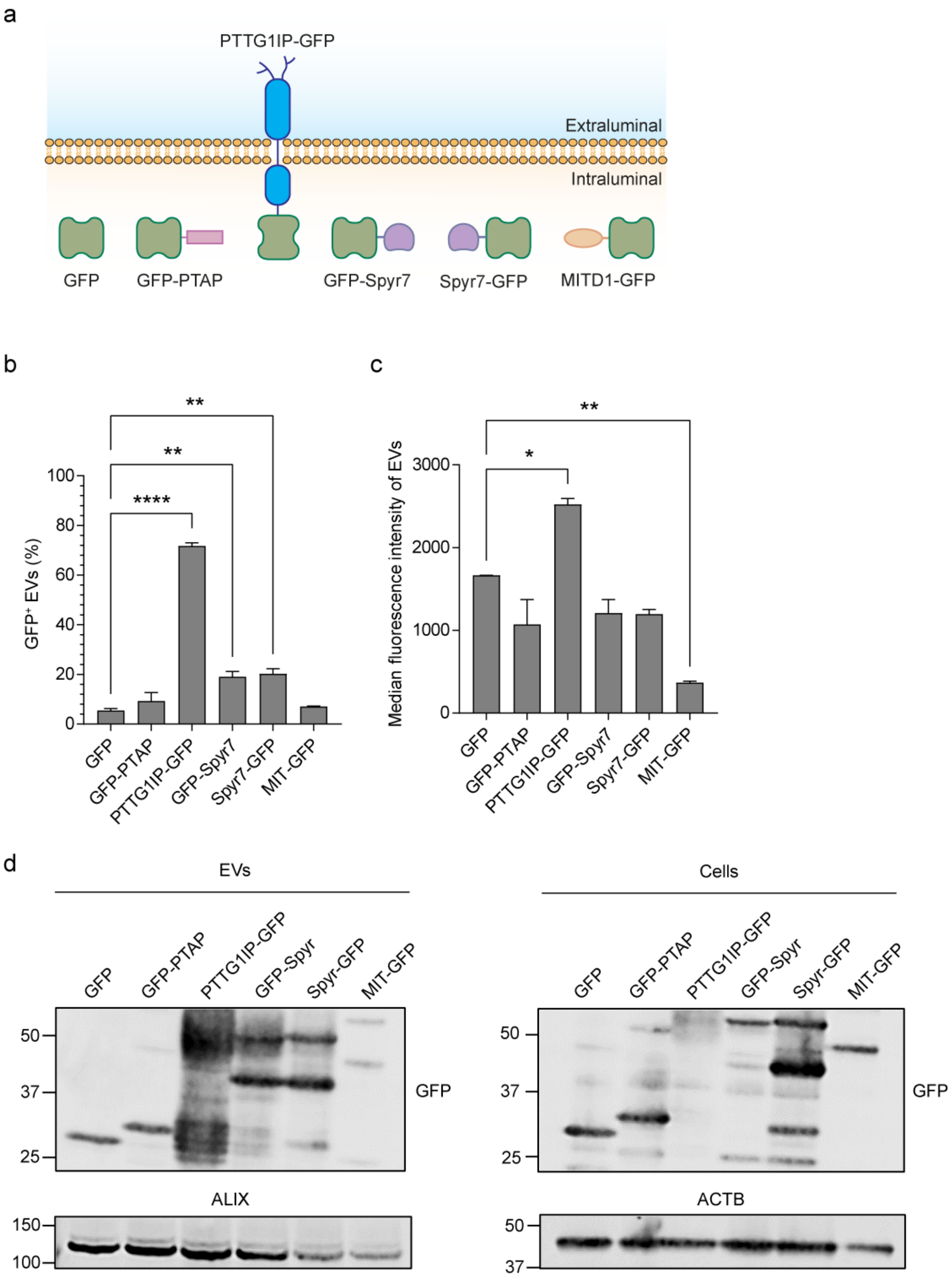
a
PTTG1IP-GFP
Extraluminal
Intraluminal
GFP
GFP-PTAP
GFP-Spyr7
Spyr7-GFP
MITD1-GFP
b
GFP⁺ EVs (%)
100
80
60
40
20
0
****
**
**
GFP
GFP-PTAP
PTTG1IP-GFP
GFP-Spyr7
Spyr7-GFP
MIT-GFP
c
Median fluorescence intensity of EVs
3000
2000
1000
0
*
**
GFP
GFP-PTAP
PTTG1IP-GFP
GFP-Spyr7
Spyr7-GFP
MIT-GFP
d
EVs
Cells
GFP
GFP-PTAP
PTTG1IP-GFP
GFP-Spyr
Spyr-GFP
MIT-GFP
50
37
25
GFP
ALIX
150
100
ACTB
50
37

**Supplementary Figure 5**

**Assessment of EV-loading features from bioinformatics analysis.**

(**a**) Scheme of the GFP-fusion constructs that were assessed. (**b**) Percentage of $GFP^+$ EVs quantified by single-vesicle analysis of conditioned media from HEK293T cells transfected with GFP-fusion constructs. (**c**) Median fluorescence intensity of $GFP^+$ EVs. (**d**) Western blot of GFP-fusion constructs in HEK293T cells and EVs. Cell lysates and isolated EVs were blotted against ACTB and ALIX respectively. Both cell lysates and isolated EVs were blotted against GFP. Statistical significance was tested by one-way ANOVA and Bonferroni *post hoc* test. Data are presented as mean + SEM, $n = 3$ replicates, $^*P < 0.05$, $^{**}P < 0.01$, $^{****}P < 0.0001$.

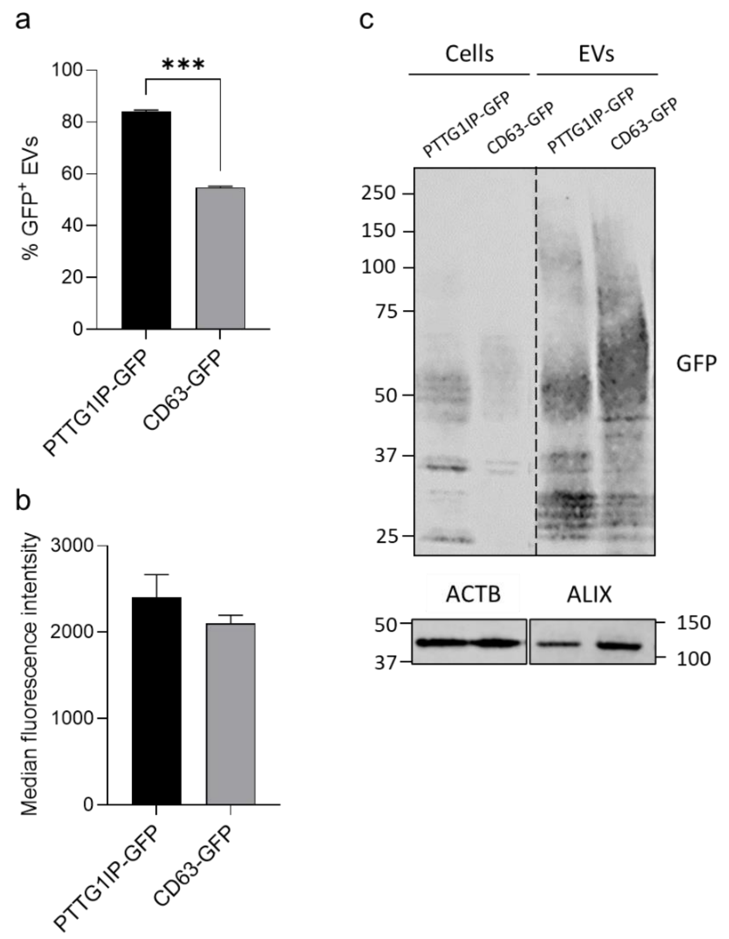


**Supplementary Figure 6**

**PTTG1IP has a higher EV-loading capacity than CD63.**

PTTGIP and CD63 EV-loading was compared by single vesicle analysis and Western Blot. (**a**) Percentage of GFP$^+$ EVs quantified by single-vesicle analysis of conditioned media from HEK293T cells transfected with PTTG1IP-GFP or CD63-GFP. (**b**) Median fluorescence intensity of GFP$^+$ EVs quantified by single-vesicle analysis. (**c**) Western blot of GFP-fusion constructs in HEK293T donor cells and EVs. Cell lysates and isolated EVs were blotted against ACTB and ALIX respectively. Both cell lysates and isolated EVs were blotted against GFP. Statistical significance was tested by Student's *t*-test. Data are presented as mean + SEM, $n$ = 3 replicates, ***$P < 0.001$.

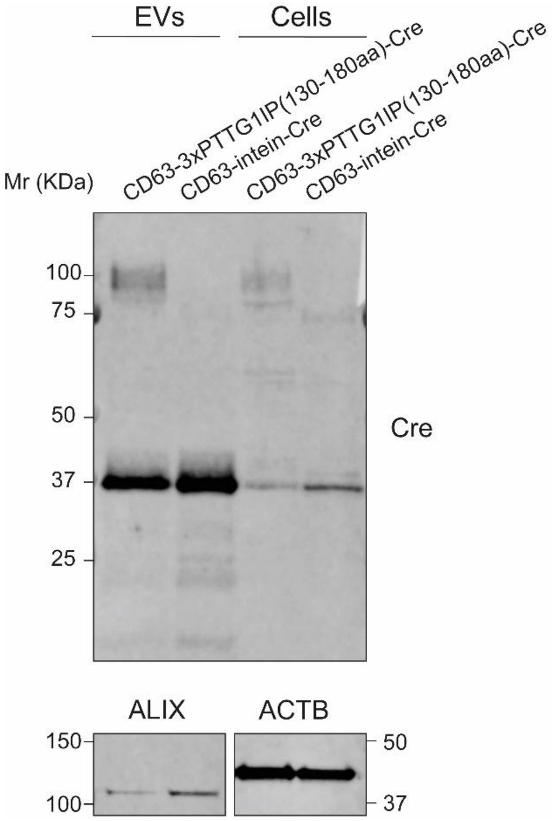


**Supplementary Figure 7**

**Amino acids 130-180 of PTTG1IP are able to induce cleavage in EVs when fused to CD63.**

HEK293T cells were transfected with CD63-intein-Cre or CD63-3xPTTG1IP(130-180aa)-Cre, and Cre cleavage in the EVs was analyzed. Western blot of CD63-intein-Cre and CD63-3xPTTG1IP(130-180aa)-Cre in HEK293T cells and EVs. Cell lysates and isolated EVs were blotted against Cre, ALIX and ACTB.

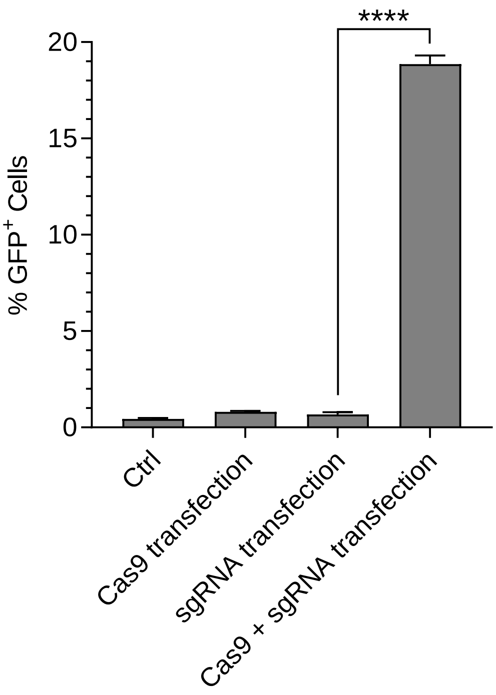


**Supplementary Figure 8**

**Assessment of Cas9 reporter cell activation after Cas9 and sgRNA transfection.**

GFP positive cells quantified by flow cytometry 48h after Cas9, sgRNA or Cas9 and sgRNA transfection to HEK293T stoplight reporter cells. Statistical significance was tested by one-way ANOVA and Bonferroni *post hoc* test. Data are presented as mean + SEM, $n = 3$ replicates, $^{****}P < 0.0001$.

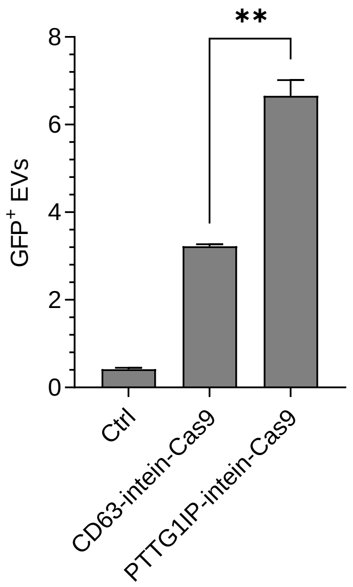


**Supplementary Figure 9**

**Comparison of PTTG1IP and CD63 for EV-mediated delivery of Cas9.**

GFP positive cells quantified by flow cytometry 48h after PTTG1IP-intein-Cas9 or CD63-intein-Cas9 EV-transfer to sgRNA transfected HEK293T stoplight reporter cells. Isolated EVs were transferred to recipient cells in 96-well plates ($2{\times}10^9$ EVs/ml). Statistical significance was tested by Student's *t*-test. Data are presented as mean + SEM, $n = 2$, $^{**}P < 0.01$.

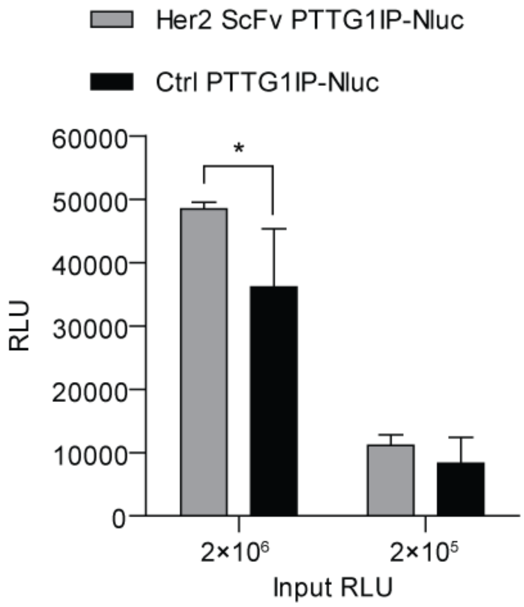


**Supplementary Figure 10**

**Uptake of targeted EVs displaying HER2 scFv in PTTG1IP N terminus by human breast cancer cell line SKBR3**.

Luciferase activity (RLU, relative light units) in recipient SKBR3 cells after treatment with EVs ($2\times10^6$ or $2\times10^5$ input RLUs) derived from HEK293T cells expressing PTTG1IP fused to HER2 scFv targeting domain at the N terminus and to Nluc at the C terminus, or EVs derived from cells expressing PTTG1IP fused to only to Nluc. Values are mean + SEM, $n$ = 3. *$P < 0.05$ (one-tailed Student's $t$-test).
